# Seasonal dynamics are the major driver of microbial diversity and composition in intensive freshwater aquaculture

**DOI:** 10.1101/2021.02.26.433039

**Authors:** Sophi Marmen, Eduard Fadeev, Ashraf Al Ashhab, Ayana Benet-Perelberg, Alon Naor, Hemant J. Patil, Eddie Cytryn, Diti Viner-Mozzini, Assaf Sukenik, Maya Lalzar, Daniel Sher

**Affiliations:** Department of Marine Biology, Leon H. Charney School of Marine Sciences, University of Haifa, Haifa, Israel; Department of Functional and Evolutionary Ecology, University of Vienna, Vienna, Austria; Microbial Metagenomics Division, The Dead Sea and Arava Science Center, Masada 86900, Israel; Dor Aquaculture Research Station, Fisheries Department, Israel Ministry of Agriculture and Rural Development, Israel; Institute of Soil, Water and Environmental Sciences, Volcani Center, Agricultural Research Organization, P.O Box 15159, Rishon Lezion, 7528809, Israel; The Yigal Allon Kinneret Limnological Laboratory, Israel Oceanographic and Limnological Research, P.O.Box 447 Migdal 14950, Israel; Bioinformatics Service Unit, University of Haifa, Israel

**Keywords:** microbiome, fishpond, 16S rRNA, cyanobacteria, runoff

## Abstract

Aquaculture facilities such as fishponds are one of the most anthropogenically impacted freshwater ecosystems. The high fish biomass reared in aquaculture is associated with an intensive input into the water of fish-feed and fish excrements. This nutrients load may affect the microbial community in the water, which in turn can impact the fish health. To determine to what extent aquaculture practices and natural seasonal cycles affect the microbial populations, we characterized the microbiome of an inter-connected aquaculture system at monthly resolution, over three years. The system comprised two fishponds, where fish are grown, and a “control” operational water reservoir in which fish are not actively stocked. Clear natural seasonal cycles of temperature and inorganic nutrients concentration, as well as recurring cyanobacterial blooms during summer, were observed in both the fishponds and the reservoir. The structure of the aquatic bacterial communities in the system, characterized using 16S rRNA sequencing, was explained primarily by the natural seasonality, whereas aquaculture-related parameters had only a minor explanatory power. However, the cyanobacterial blooms were characterized by different cyanobacterial clades dominating at each fishpond, possibly in response to distinct nitrogen and phosphate ratios. In turn, nutrient ratios may have been by the magnitude of fish feed input. Taken together, our results show that, even in strongly anthropogenically impacted aquatic ecosystems, the structure of bacterial communities is mainly driven by the natural seasonality, with more subtle effects if aquaculture-related factors.

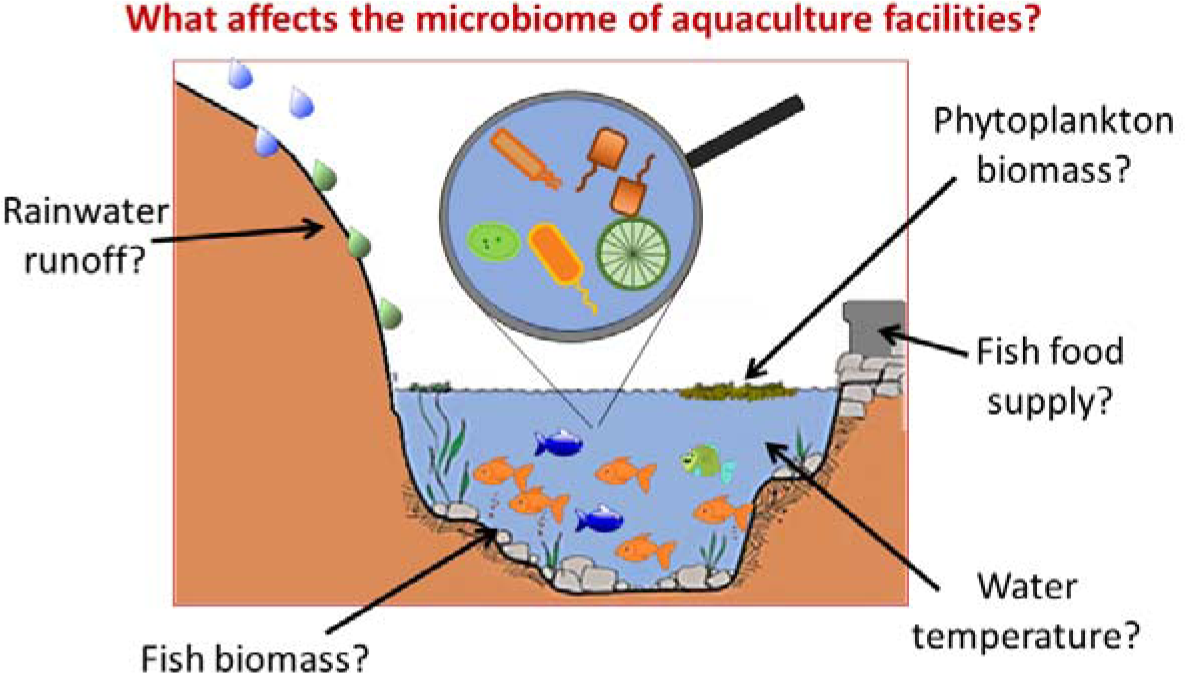

**Highlights:**

- We present three years of monthly microbiome data from an aquaculture facility.
- The microbiome changes seasonally, likely driven by temperature and rainwater runoff.
- Summer blooms of toxin-producing cyanobacteria are repeatedly observed.
- Fish food may impact microbiome through changes in nutrient ratios.

## 1. Introduction

Freshwater environments come in many shapes and forms, from pristine mountain lakes to highly polluted wastewater treatment facilities, each hosting a complex and dynamic microbial community (“microbiome”; Bautista-de los Santos et al., 2019; Bruno et al., 2018; Zorz et al., 2019). Such aquatic microbiomes are strongly affected by environmental conditions within the water body (e.g., temperature and nutrients availability), and in turn shape its biogeochemistry, for example by processing and recycling organic and inorganic matter (Hach et al., 2020; Liu et al., 2017; Rowe et al., 2018). Aquatic microbial communities may also affect other organisms living within the water body, as well as human communities in contact with the water. For example, the eggs and larvae of some aquatic organisms are inoculated by microbes from the surrounding water, in turn affecting the health and fitness of these organisms (Giatsis et al., 2016; Grze kowiak et al., 2012; Pérez-Sánchez et al., 2014; Song et al., 2014). Similarly, aquatic microbiomes may serve as pools of various pathogens, and some aquatic microorganisms (primarily prokaryotic and eukaryotic microalgae) can produce toxins (Paerl et al., 2018; Wurtsbaugh et al., 2019). Given the importance of freshwater microbiomes to water quality and environmental health worldwide, significant effort has been invested in understanding the interplay between environmental conditions and microbiome structure and function (reviewed by Orland et al., 2019).

Freshwater aquaculture facilities such as fishponds are an example of an aquatic ecosystem that is significantly impacted by anthropogenic activity. The growth of human population and the concomitant increase in the demand for animal-based protein have resulted in a rapid expansion of aquaculture worldwide, a process also termed a “blue revolution” (reviewed by Garlock et al., 2020). The aquaculture industry has been growing at an average rate of ca. 6-8% since the 1970s, and now supplies more fish and seafood biomass than natural catch (Ahmed and Thompson, 2019; Garlock et al., 2020). As a result, fishponds (as an ecosystem) are becoming more abundant, and their interaction with surrounding environments is increasing.

Modern intensive aquaculture practices rely on growing large stocks (i.e., high biomass) of fish or other aquatic organisms in a limited area. This requires large amounts of fish-feed that often leads (together with fish excrement) to hyper-eutrophic conditions (Ahmed and Thompson, 2019; Gowen, 1994; Talbot and Hole, 1994). Additionally, aquaculture is often performed in monoculture (i.e., a single type of organism is grown) or in polyculture of a limited number of organisms, and the combination of ultra-dense populations and hyper-eutrophic conditions (i.e., poor water quality) can lead to disease outbreaks (Ahmed and Thompson, 2019; Glibert et al., 2002; Mennerat et al., 2010). As a result, aquaculture often supports fish health by using drugs and antibiotics, as well as algaecides in the case of algal blooms. However, despite the ubiquity of such aquaculture husbandry practices, and their potential impact on aquatic microbial populations, little is known about the temporal dynamics of the microbiome in fishponds, and how it responds to natural and anthropogenic perturbations (Patil et al., 2020; Rodriguez-Brito et al., 2010).

To address these questions, we characterized, over three years, temporal microbiome dynamics in two fishponds connected to a semi-natural operational water reservoir, which recycles the fishpond water (Fig. 1, see detailed description below). Our aim was to identify factors that dictate the water microbiome composition of these highly anthropogenically impacted ecosystems. These included the quality of the water, phytoplankton populations, as well as to the types of fish present and their feeding regimen. We hypothesized that specific intensive fishery practices in each fishpond (e.g., the types of fish raised and their feeding regime) would shape the microbiomes of the fishponds, resulting in different populations between the fishponds, as well as between them and the reservoir. Our results, however, point to seasonality rather than specific aquaculture practices as the major force driving the microbial population structure.

**Fig. 1.**
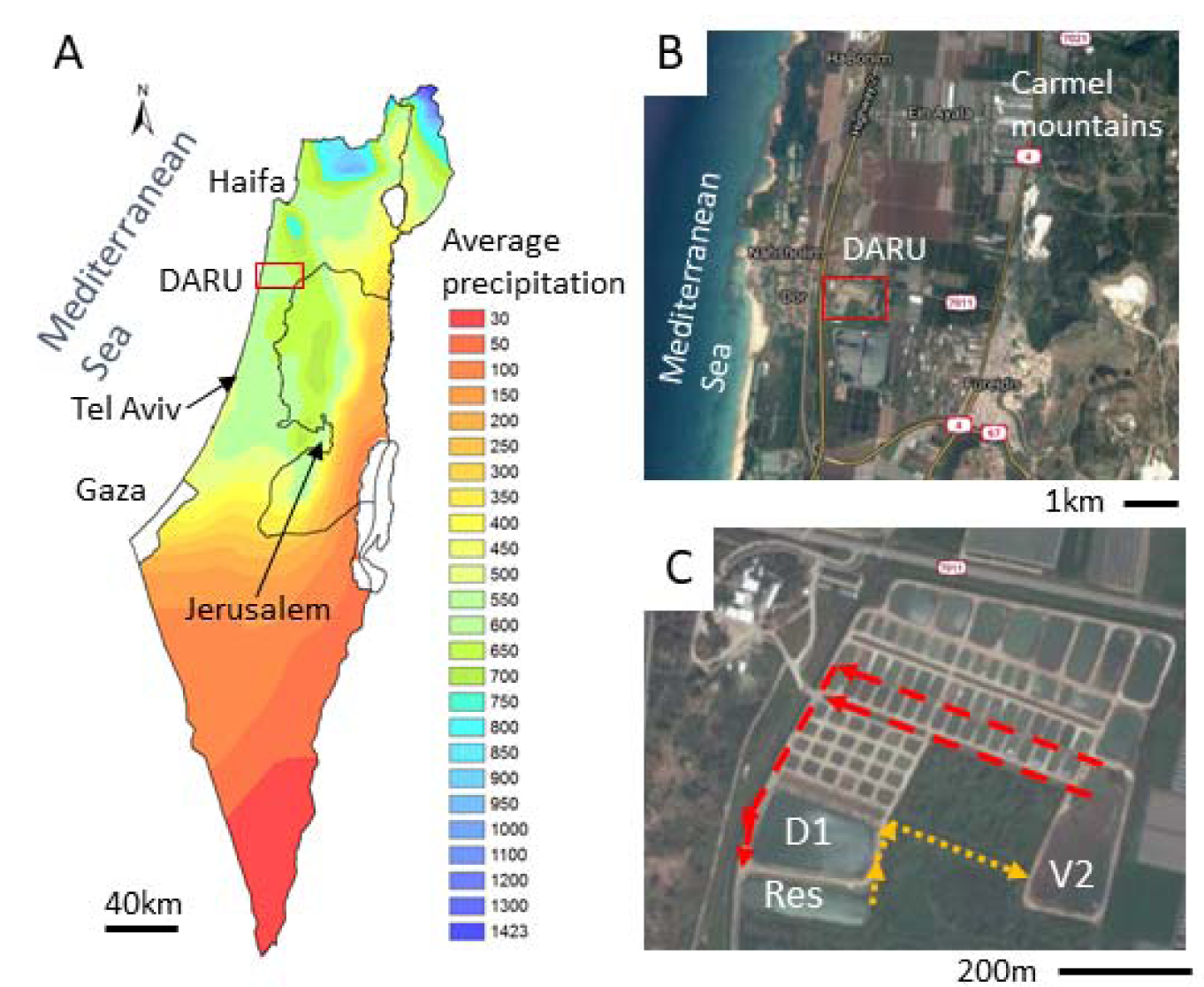
The Dor Aquaculture Research Unit (DARU). a) Mean annual precipitation map of Israel, showing DARU on the coastal region which has a Mediterranean climate. The red square marks the location of panel b. b) The coastal plain between the Carmel mountains and the sea, with the location of DARU (panel c) shown as a red square. DARU is situated in a predominantly agricultural region with banana plantations and various fields. c) DARU. The arrows show the flow direction between the D1 and V2 fishponds and the Reservoir. The mean annual precipitation map of Israel was derived from the digital elevation model (DEM) using ArcGIS (ESRI, Redlands, CA, USA).

## 2. Materials and Methods

### 2.1 Sampling and analysis of environmental data

Monthly samples were collected at the reservoir and fishponds D1 and V2 (when they contained water) from 2013-2015 (Table S1). Each pond was sampled at a constant location from the edge of the water body and included measurement of dissolved oxygen, temperature and pH, using field probes (Eutech instruments, Singapore). At each location, five liters of surface water were collected using plastic jerrycans (specific to each location) that were washed with double distilled water and with 70% ethanol prior to sampling. Within one hour the samples were transferred in darkness to the lab and processed according to the methodology described below.

For measurement of pigments, water was filtered through GF/C filters (glass fiber filter, 13 mm, nominal pore size 1.2 μ; Whatman plc, Maidstone, UK) and placed into 1.5 mL sterile Eppendorf tubes. The water was filtered until the filters were clogged, and the volume was recorded (30-150 mL). For DNA extraction and microcystin measurements, water was filtered through GF/F filters (glass fiber filter, 13 mm, nominal pore size 0.7 μm; Whatman plc, Maidstone, UK). The filters were placed into 1.5 mL sterile tubes, and for the purpose of DNA extraction were covered with 1 mL of storage buffer (40 mM EDTA, 50 mM Tris-HCl, 0.75 M Sucrose) and stored at -80 °C until analyzed. The filtrate from the GF/F filters was collected for measurement of dissolved inorganic nutrients, and were maintained at -20 °C until analysis.

### 2.2 Inorganic nutrients measurements

The concentrations measurements of PO_4_^+^, NH_4_^+^, NO_3_^-^ and NO_2_^-^ in fishponds and in the water reservoir were measured using the AA3 Segmented Flow Multi-Chemistry Analyzer (SEAL Analytical, Germany), following the manufacturer protocols (Ammonia: Method no. G-327-05 Rev. 7 -Fluorescent method; Nitrate and Nitrite: Method no. G-172-96 Rev. 17 and Phosphate: Method no. G-297-03 Rev. 5). Inorganic nutrients in the rainwater runoff were measured using a Lachat Autoanalyzer (Lachat Instruments, Milwaukee, WI, USA) by the manufacturer’s protocol.

### 2.3 Photosynthetic pigment analysis

Pigments were extracted for 3 hours in absolute methanol in the dark. Pigment extractions were immediately clarified with syringe filters (Acrodisc CR, 13 mm, 0.2 µm PTFE membranes; Pall Life Sciences, New York, NY, USA) and transferred to glass ultra-performance liquid chromatography (UPLC) vials. The samples were preheated to 30 °C, and 10 µl were injected into an ACQUITY UPLC system (Waters Corporation, Milford, MA, USA) equipped with a C8 column (1.7 µm particle size, 2.1 mm internal diameter, 50 mm column length, ACQUITY UPLC BEH) heated to 50 °C and a guard column. The protocol was based on the LOV method (Hooker and McClain, 2000), with some modifications: the mobile phase gradient consisted of a mixture of 70:30 methanol: 0.5M ammonium acetate as solvent A and 100% methanol as solvent B. Peaks were monitored at 440 nm and their absorbance spectra was determined using a photodiode array (PDA) detector. Known standards of Chlorophyll a, Chlorophyll b, Chlorophyll c_2_, Zeaxanthin, β-carotene, Diatoxanthin, Dinoxanthin, Fucoxanthin and Peridinin were separated before each run for further identification and quantification. All standards were purchased from the DHI Laboratory (Hørsholm Denmark). Pigments were identified by retention time and spectrum absorbance (obtained by PDA detector reads at 350-700 nm). Pigment concentrations were adjusted to 1 mL extraction volume and filtration volume.

### 2.4 Fish biomass and supplied food mass estimation

Fish biomass and the fish-feed input were estimated for the sampling time-points based on the limited records kept at the Dor Aquaculture Research Unit (DARU). When the fishponds were established for the growing season, the total number and average biomass of the larvae or juvenile fish was recorded. Once or twice a month, during the outgrowth period, a representative sample of fish was weighed and returned to the water. The number of fish in the water was estimated and the amount of fish-feed supplied was recorded. Over the course of the last several weeks of the growing period, fish are often harvested in batches, resulting in a gradual decrease in the number of fish in the fishpond. During each harvesting event, a representative sample of the removed fish was weighed. At the end of the growing period (as the water is removed and the fishponds dried until the next growing season), the final number of fish and their weight were recorded. Based on these measurements, we calculated the fish biomass and used a fitted model in Microsoft Excel (Table S2, tabs “D1_calculations“ and V2_calculations”) to estimate the fish biomass when our samples were taken, which was always during the outgrowth period. A similar approach was applied to estimate the amount of food added to the fishponds. A second order polynomial model provided a good fit to most of the data (R^2^ above 0.96; Table S2, tabs “D1_calculations“ and V2_calculations”). We would like to emphasize that these models are not designed to be predictive or mechanistic and are limited to interpolating the data to provide estimates of the biomass and feed mass during the sampling times.

### 2.5 Microcystin extraction and measurement

Cyanotoxins (microcystins or nodularins) in phytoplankton biomass were measured immunologically. Biomass collected on filters was subjected to methanolic extraction (3 hours incubation in 1 mL 100% Methanol), followed by 15 minutes sonication. The upper 500 µL were dried under vacuum for 2 hours in a speedvac, resuspended in 50 µL of 75% methanol and further dilution to 4% methanol. A subsample of 50 µL of that solution was used for quantification by the Microcystins/Nodularins (ADDA) Elisa Kit (Eurofins Abraxis Inc, Warminster, PA, USA). This protocol led to a concentration factor of 5-100 (depending on the initial filtered volume), bringing the samples within the range of sensitivity of the measuring kit (0.1 µg L^-1^). In some cases, the ELISA results t were above the linear range of the kit, therefore, these samples (marked in Fig. 2, Fig. S1, and Table S2) should be considered as likely underestimates.

**Fig. 2.**
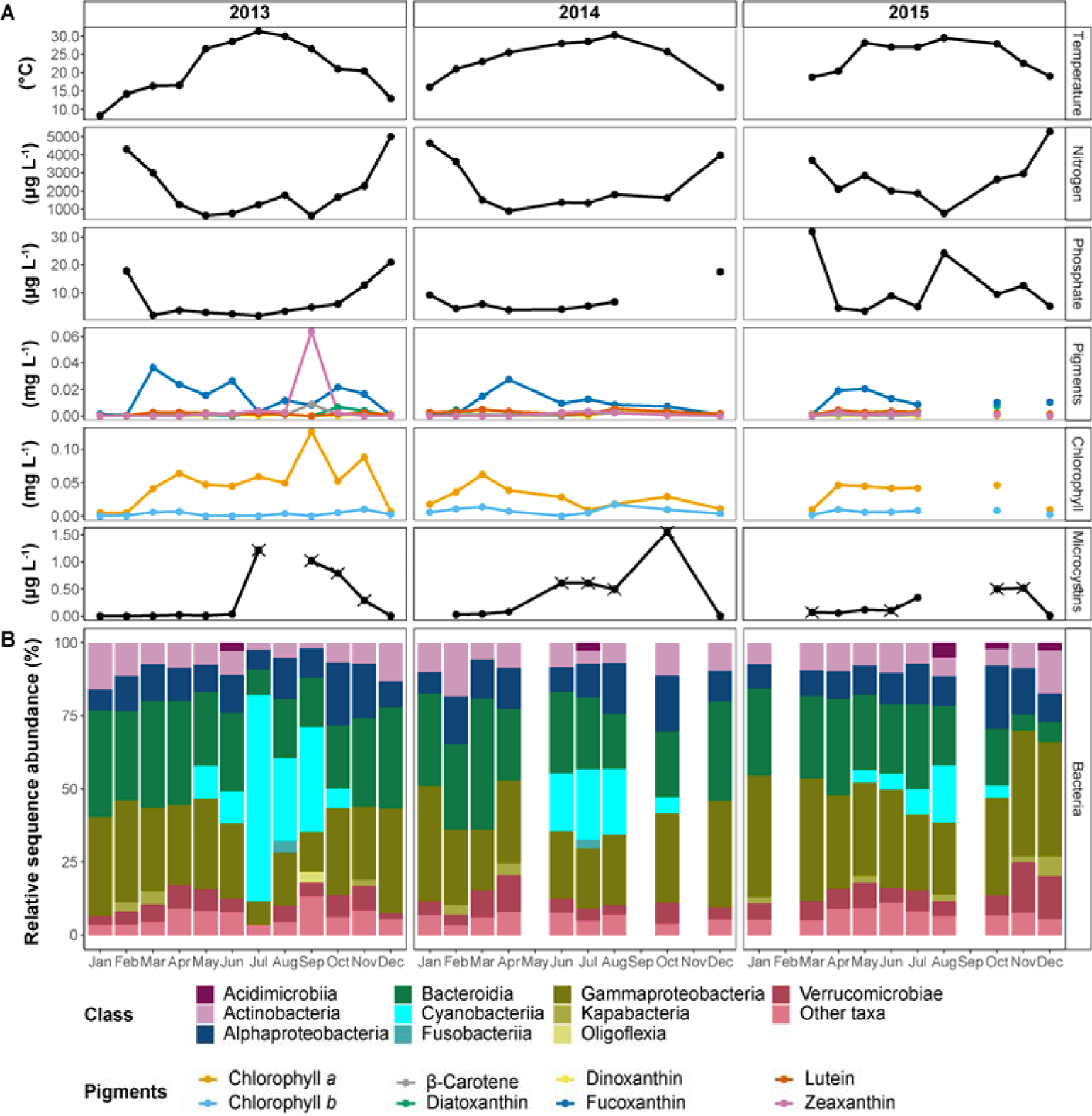
Seasonal dynamics in the Reservoir between 2013 and 2015. (a) Monthly measured physicochemical properties of the water. The different photosynthetic pigments are colored according to the legend. Nitrogen represents the total concentration of nitrate and nitrite. In microcysteins, samples marked with ‘×’ represent potentially underestimated concentration. For visualization purposes, microcysteins measurements of Oct. 2014 and Aug. 2015 (1.6 and 10.4 µg L^-1^, respectively) were omitted from the figure. (b) Sequence proportion overview of bacterial communities on a class level. The classes represented by colors according to the legend, all classes with sequence proportions below 2% were classified as “Other classes”.

### 2.6 DNA isolation and 16S rRNA amplicon sequencing

The DNA extraction was performed using a combination of manual treatments and robotic extraction with the QiaCube robot (Biomedical Core Facility, Technion, Haifa, Israel). Briefly, filters were thawed, centrifuged for 10 minutes at 15,000 ×*g* and the storage buffer was removed. Lysis buffer from the DNeasy Blood & Tissue Kit (Qiagen, Hilden, Germany) was added and the samples were mechanically pounded with two 3 mm sterile stainless steel beads at the speed of 30 Hz for 1.5 minutes using a TissueLyser LT (Qiagen, Hilden, Germany). After addition of 30 µl lysozyme the tubes were incubated at 37 °C for 30 minutes, followed by the addition of 25 mL proteinase K and 200 µL of the buffer AL with an additional incubation for 1 hour at 56 °C on a shaker. Finally, the tubes were centrifuged for 10 minutes at 5,000 ×*g* and the upper liquid was transferred to a new 2 mL tube for extraction by the QiaCube robot, according to manufacturer’s instructions (Qiagen, Hilden, Germany). The DNA quantity was assessed using Picogreen (Quant-iT PicoGreen dsDNA reagent; Invitrogen, Carlsbad, CA, USA), and the quality (apparent size and degradation products) was assessed using a TapeStation (Agilent Technologies, Santa Clara, CA, USA).

Polymerase chain reaction (PCR) reactions for amplifying 16S rRNA gene fragments were performed in triplicate for each DNA sample, using the following primers that target the V3-V4 region: forward CS1_341F (Muyzer et al., 1993) (5’-ACA CTG ACG ACA TGG TTC TAC ANN NNC CTA CGG GAG GCA GCA G-3’) and reverse CS3_806R (Caporaso et al., 2011) (5’-TAC GGT AGC AGA GAC TTG GTC TGG ACT ACH VGG GTW TCT AAT-3’). The PCR reactions were performed in a final volume of 25 μl, with 10 ng of DNA template. The PCR protocol was as follows: initial denaturation stage at 95 °C for 5 minutes, followed by 28 cycles at 95°C for 30 seconds, 50 °C for 30 seconds, and 72 °C for 60 seconds, with a final extension step at 72 °C for 5 minutes, using BIOLINE 2x MyTaq Red Mix (Sigma-Aldrich, St. Louis, MO, USA), and the PCR reaction was performed in a TProfessional Basic Gradient thermocycler (Biometra, Göttingen, Germany). Following the first PCR, triplicate samples were pooled, and the products were sent to the DNA Services Facility of the University of Illinois, Chicago, where a second PCR was performed to incorporate barcodes and sequencing adapters (CS1 and CS3). Sequencing was performed using a 2×250 base pair format on a MiSeq flow cell (V3 chemistry, Illumina, USA). The 2013-2014 and 2015 samples were sequenced at different times and on two separate sequencing lanes (see results). The raw sequencing reads are available as BioProject PRJNA688121 at the NCBI SRA database.

### 2.7 Bioinformatics and statistical analyses

The raw paired-end reads were primer-trimmed using cutadapt (Martin, 2011), and subsequent analyses of the generated sequences were conducted using R (v3.6.3; http://www.Rproject.org/) in RStudio (v1.2.5033; http://www.rstudio.com/) as described below. The trimmed libraries were processed using DADA2 (Callahan et al., 2016) (v1.14.1), following the suggested tutorial (https://benjjneb.github.io/dada2/tutorial.html). Initially, chimeras and singletons were filtered out, and subsequently, amplicon sequence variants (ASVs) were taxonomically classified against the Silva reference database (release 138) (Quast et al., 2012). The ASVs that were taxonomically unclassified to the domain level, or not assigned to bacterial or archaeal lineages, were excluded from further analysis. Furthermore, all ASVs which were taxonomically assigned to mitochondria and chloroplast were removed from the dataset.

Sample data matrices were managed using the R package ‘phyloseq’ (v1.28.0; McMurdie and Holmes, 2013) and plots were generated using R package ‘ggplot2’ (v3.3.0; Wickham, 2016). Calculation and visualization of shared ASVs was conducted using R package ‘venn’ (v1.9; Dusa, 2020). Statistical tests were conducted using R package ‘vegan’ (v2.5.6; Oksanen et al., 2019). The sample rarefaction analyses were conducted using R package ‘iNEXT’ (v2.0.20; Hsieh et al., 2019). All beta-diversity and RDA analyses were conducted on a transformed ASV table using a geometric mean. The fold-change in abundance of each ASV between the water layers was calculated using the R package ‘DEseq2’ (v1.24.0; Love et al., 2014). The method applies a generalized exact binomial test on variance stabilized ASV abundance. The results were filtered by significance, after correction for multiple-testing according to Benjamini and Hochberg (1995) with an adjusted p-value <0.1.

Scripts for the molecular data processing and statistical analyses can be accessed at https://github.com/edfadeev/Dor_ponds_16S

## 3. Results

### 3.1 Description of the study site

The Dor Aquaculture Research Unit (DARU - 32°36′25.4 N 34°55′54.7″E) is situated on the coastal plain of Israel (Fig. 1). DARU belongs to the Ministry of Agriculture and Rural development of Israel and operates as a research unit and a semi-commercial outgrowth facility. The research at DARU is performed in approximately 80 miniature ponds (0.03 hectare), and focuses on genetically-informed breeding programs, increasing fish health by testing new immunization methods and developing healthier and more ecologically-sound methods of intensive fish rearing (Patil et al., 2016). In addition, DARU has two larger ponds, termed D1 and V2 (1.8 and 1.6 hectares each, with 18,000 and 12,000 m^3^ of water, respectively), which are used for semi-commercial outgrowth of Common carp (*Cyprinus carpio*) and Silver carp (*Hypophthalmichthys molitrix*). All the ponds are connected to an external, one-hectare, operational reservoir (ca. 8,000 m^3^), which is used to regulate the water levels and temperature in the fishponds. Water is supplied to the reservoir from a local well and from floodwater during the winter months. The ponds and the reservoir are connected by a series of channels (Fig. 1). During the three-year study period, fishpond D1 was stocked with Common carp, whereas fishpond V2 was stocked with Silver carp in 2013 and with Common carp in 2014-2015 (Table S1). The outgrowing period of the Common carp is typically about 5-7 months whereas that of a Silver carp is shorter (ca. 2-3 months), both occurring during the spring and summer months (i.e., March-September). When no fish are grown the fishponds are kept dry, with the exception of occasional flood-water puddles at the deepest part of the fishpond. At the initiation of the outgrowth period, two-thirds of the fishponds’ volume is filled with water pumped from the operational reservoir, and one-third with fresh groundwater from a local well. During the growing period, the water temperature is regulated by adding cold water from the well, or warmer water from the reservoir. At the end of the growing period, water from the fishponds is transferred through connecting channels to the operational reservoir, and then via a natural stream to the Mediterranean Sea. The reservoir contains water throughout the entire year, and was sampled almost monthly, and primarily during the outgrowth period in fishponds D1 and V2.

### 3.2 Water quality parameters change seasonally in the reservoir

We first focus on the operational reservoir, which contains water throughout the entire year and provides a baseline for the system over the three years of sampling (Fig. 2). As expected from a water body situated in a typical Mediterranean climate (cold and wet winters and hot and dry summers), and despite the capacity at DARU to regulate water temperature to some extent by adding well-water, the temperature showed a clear seasonal cycle, ranging from 10-20 □ during the rainy season (November-April; further addressed as ‘wet season’) to 20-30 □ during the dry season (May-October; further addressed as ‘dry season’; Fig. 2a). Inorganic nutrient concentrations were typically low during the summer months, began rising in autumn and reached the highest concentrations during the winter months, although some variability was observed from year to year (Fig. 2a). Importantly, we had expected that the nutrients in the reservoir would be higher during the fish outgrowth period (spring and summer), when intensive fish feeding occurs, as a potential result of water from the fishponds reaching the reservoir (Fig. S1a). The observation that nutrient concentrations peaked during winter prompted us to search for other, non-aquaculture related, processes that might have provided nutrient input to the reservoir. One such process is rainwater runoff from agricultural fields and banana plantations surrounding DARU (Fig. 1b). Indeed, runoff water sampled during one rain event (Feb 12, 2015) had high concentrations of inorganic nutrients (270-390, 90-140 and 1200-5600 µg L^-1^ of ammonia, nitrite and nitrate, respectively). These concentrations are within the range of the highest concentrations measured in the reservoir over the time-series.

### 3.3 Water quality parameters, fish biomass and feeding differed between the two fishponds

Similar to the reservoir, the temperature changed seasonally in the two fishponds, D1 and V2, and dissolved inorganic nitrogen concentrations were typically higher during the winter months (even when fish had not been stocked yet; Fig. S1). However, there were major differences between the fishponds, and between them and the reservoir. The reservoir waters consisted of up to tenfold higher concentrations of inorganic nitrogen (N; 2017±263 µg L^-1^, n=23), compared to the fishponds D1 (619±301 µg L^-1^, n=19) and V2 (464±222 µg L^-1^, n=20; Kruskal-Wallis test, Chi square=17.61, df=1, p<0.001). On the other hand, both fishponds reached up to tenfold higher concentrations of soluble reactive phosphate (P; D1 - 59±25 µg L^-1^, n=17; V2 - 91±21 µg L^-1^, n=19), compared to the reservoir (8±2 µg L^-1^, n=22; Kruskal-Wallis test, Chi square=17.66, df=1, p<0.001). The N:P ratio also strongly differed between the ponds. In both the reservoir and the fishpond D1 the median N:P ratio (i.e., the ratio in half of the sampled time points) was equal or above the Redfield ratio of 16N:1P (ca. 275 and ca. 16, respectively). In contrast, in fishpond V2 the median N:P ratio was close to 1. This similarity pattern between the water bodies was also observed in the chlorophyll *a* (chl-*a*) concentrations. In the reservoir and in fishpond D1, the chl-*a* was lower (0.04±0.01 mg L^-1^; and 0.03±0.01 mg L^-1^, n=29, n=16 respectively) compared to the fishpond V2 (0.12±0.03 mg L^-1^).

Over the three years of the study, the fish biomass differed between the two fishponds, D1 and V2 (Fig. S2; see materials and methods for the estimation of biomass and feed from routine aquaculture measurements). During the times where fish were maintained in both ponds, the total estimated fish biomass was always higher in fishpond D1 than at fishpond V2 (Fig. S2). However, the estimated feed input was similar between the two fishponds. Some fish also live in the reservoir, however, they are not actively stocked nor provided with food, and thus the biomass there was assumed to be negligible compared to the densely populated fishponds.

### 3.4 Seasonal dynamics are observed in reservoir and the fishponds microbiome structures

To obtain an overview of the microbial populations in the reservoir and the fishponds, we sequenced 16S rRNA gene amplicons using Illumina technology. The final dataset consisted of 1,840,160 sequences from 74 samples that were assigned to 5,236 ASVs Amplicon Sequence Variants (Table S2). Rarefaction curves reached a plateau in all of the sampled communities (Fig. S3). An estimated asymptotic extrapolation of the curves to double the amount of sequences (i.e., theoretical deeper sequencing) showed only few additional ASVs, thus suggesting that our sequencing effort was sufficient to represent most of the bacterial diversity. Although identical extraction and amplification protocols were used, we observed a strong difference in the number of sequences per sample and the number of observed ASVs between 2013-2014 and 2015, which were sequenced separately (Fig. S3). Therefore, despite the overall similarity in community composition at high taxonomic classifications (e.g., class; Fig. 2b and Fig. S4), further statistical analyses were mainly performed on the 2013-2014 samples, with separated complementary analysis of the 2015 samples.

Overall, the bacterial communities of all three water bodies were strongly dominated by the classes *Actinobacteria*, *Alphaproteobacteria*, *Bacteroidia*, *Gammaproteobacteria*, and a prominent bloom of *Cyanobacteria* was observed during the summer months (Fig. 2 and Fig. S1). Richness and diversity indices did not show significant differences between the bacterial communities of the different water bodies (Table S2 tab “Alpha_diversity“; Kruskal-Wallis test, *p*>0.05). The communities of the reservoir and the fishponds shared 497 ASVs (one-fifth of total observed ASVs; Fig. S4), that comprised ca. 30-80% of the sequences in the bacterial communities. Each water body contained an additional ca. 500 unique ASVs (one third of the ASVs observed in each water body), which constituted up to 15% of the total ASVs. The rest of the ASVs were shared between two of the three water bodies.

We had expected to observe major differences between the fishponds, where high fish biomass is grown, and the reservoir, in which fish are not actively stocked. Indeed, significant differences in the bacterial community composition of the two water bodies were revealed (Fig. S5a; PERMANOVA test, *F_2,42_*=1.55, R^2^=0.06, *p*<0.01), however, seasonal effects on the microbiome appeared to be a stronger explanatory variable (PERMANOVA test, *F_1,42_*=5.5, R^2^=0.10, *p*<0.001). This suggests similar seasonal dynamics of the microbial population in all three water bodies. Furthermore, a separate analysis of the bacterial communities in 2015 revealed similar pattern of a significant dissimilarity between the water bodies (PERMANOVA test, *F_2,20_*=1.66,R^2^=0.12, *p*<0.01), which was lower than between the seasons (PERMANOVA test, *F_2,20_*=2.45, R^2^=0.09, *p*<0.001).

### 3.5 Specific ASVs enriched in each water body

Using comparative sequence enrichment tests, we identified the bacterial genera that had significantly different relative sequence abundance between wet and dry seasons in all water bodies combined. The ASVs within each genus were defined as enriched when they had a log_2_ fold change >1 and an adjusted *p* value <0.1. We found that 84 and 74 ASVs were significantly enriched in the bacterial communities of dry and wet seasons, respectively (Fig. 3). In both seasons the sequences of these enriched ASVs comprised 20-60% of the bacterial communities (Fig. 3). In the wet season, the enriched ASVs of the genera *Limnohabitans* (class *Gammaproteobacteria*; total of 11 ASVs) comprised 8±1% of the bacterial communities. In the dry season, a quarter of the enriched ASV (total of 23) were of *Cyanobacteria*. Enriched ASVs of the cyanobacterial genera *Cyanobium* (15 ASVs) comprised 8±1% of the bacterial communities. Furthermore, there were 2 significantly enriched ASVs from the genus *Microcystis*, which comprised 5 ± 2% of the bacterial communities. Thus, the cyanobacterial blooms are a pervasive phenomenon underlying the differences in community composition between the wet and dry seasons. These blooms were associated with the presence of cyanobacterial toxins in the water (microcystins or nodularins, Fig. 2a and Fig. S1).

**Fig. 3.**
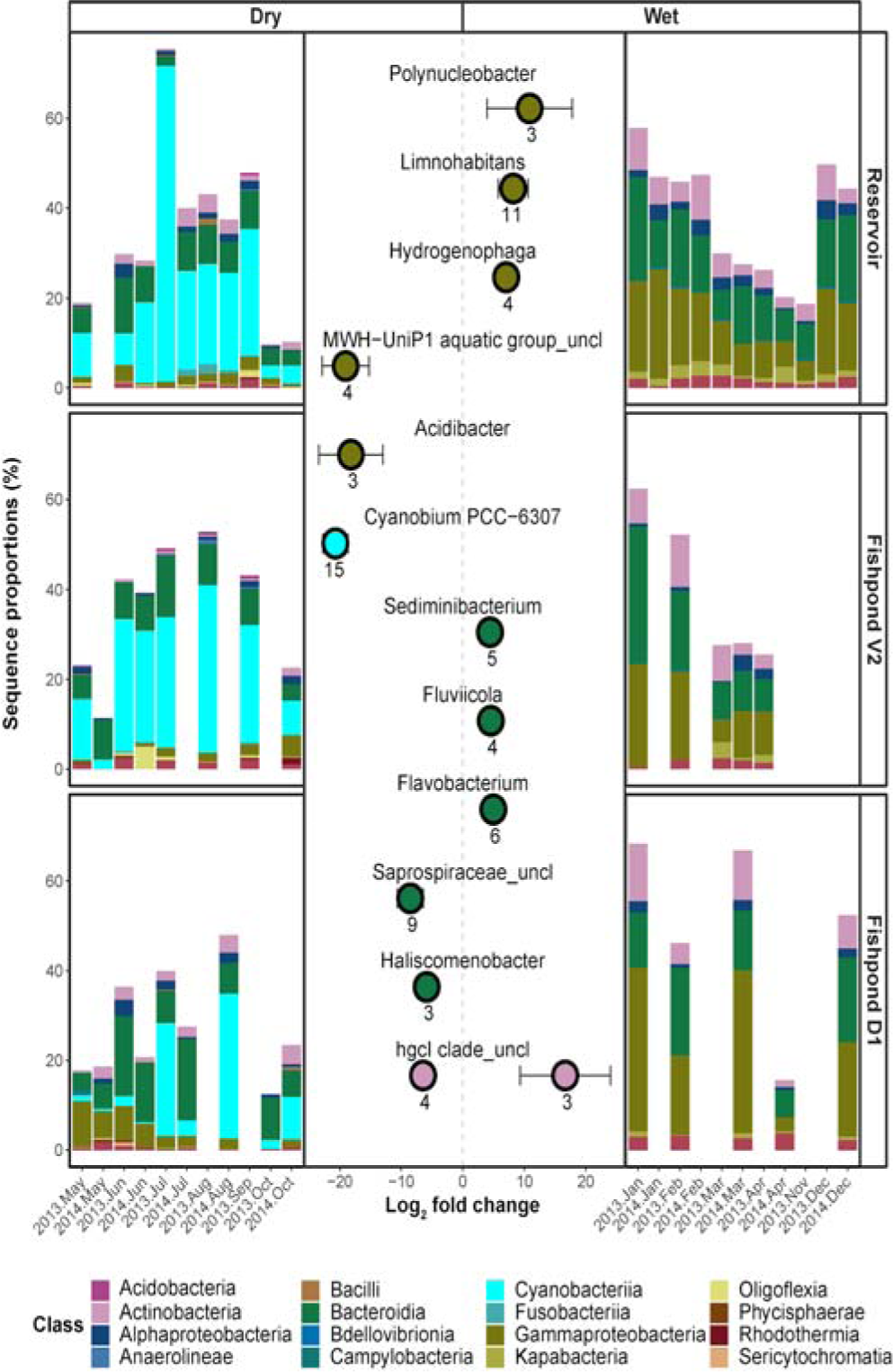
Differential enrichment analysis comparing ASV sequence abundances during dry season (left) and wet season (right). The x-axis of the central plot represents the log_2_ fold change for all significantly enriched ASVs within each labeled genera. Positive and negative values represent enrichment during wet and dry seasons, respectively. Each point and bars represents the mean and the standard deviation of the ASVs log_2_ fold change in each genera. Only taxonomic families with at least two significantly enriched (adjusted p<0.1) ASVs, and an absolute log_2_ fold change value of 1, were included in the figure. The peripheral barplots represent the respective sequence proportions of the enriched ASVs within the bacterial communities of the different pools.

### 3.6 The effect of physicochemical conditions on the bacterial community dynamics

In order to identify whether the observed seasonal changes in the bacterial communities are associated with natural physicochemical dynamics or with aquaculture-related dynamics we performed redundancy analyses (RDA) constrained by various measured and estimated parameters. Temperature, inorganic nutrient concentrations (phosphate, ammonia, nitrate and nitrite), as well as N:P ratio and microcystin concentrations, showed the highest explanatory power, explaining 15% of the total variation in the bacterial communities. These parameters separated the bacterial communities of wet and dry seasons, in accordance with the unconstrained dissimilarity analysis (Fig. 4a). The wet season bacterial communities were associated with lower water temperatures and higher concentrations of N, whereas the bacterial communities of the dry season were associated with higher temperatures and to some extent with higher microcystin concentrations.

**Fig. 4.**
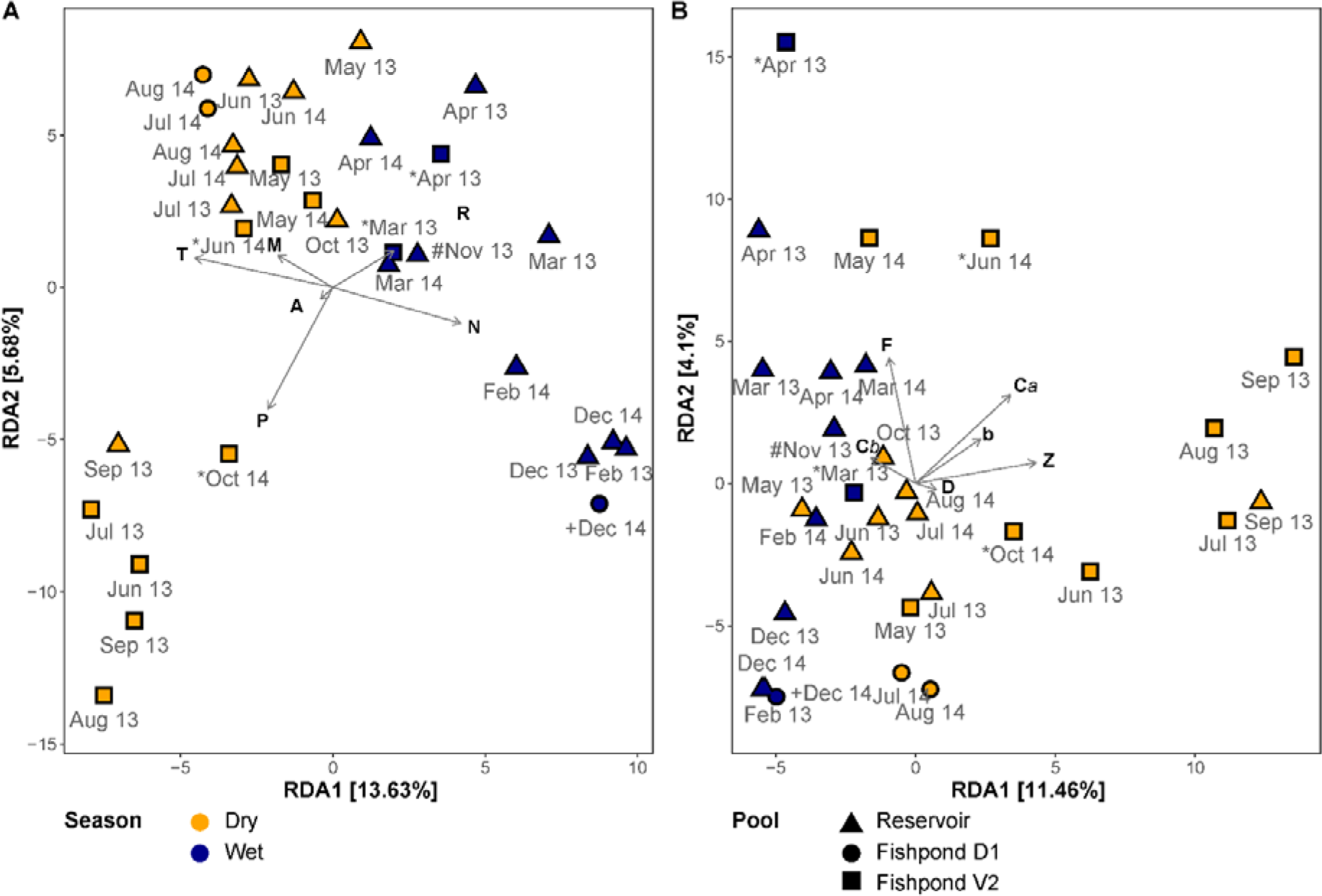
RDA ordination of bacterial community composition constrained by physicochemical parameters (a) and photosynthetic pigments (b). Colors represent the seasons (dry – April to October; Wet –November to March) and shapes represent the different ponds. The percentage on the X and the Y axis represent the proportion of explained variance. The environmental variables are: T - temperature, N - inorganic nitrogen, P - inorganic phosphate, A - ammonia, M -microcystins, R - the N:P ratio The pigments are: C*a*- chlorophyll *a*, C*b*- chlorophyll *b*, b - β-carotene, D - Dinoxanthin, F - fucoxanthin, Z - Zeaxanthin. Special conditions: (*) – no fish in the fishpond, (#) – decomposed corpse of a donkey at the edge of the reservoir, (+) – winter puddle in a fishpond, outside of the culturing season.

Interestingly, in 2013 we observed association between the bacterial communities of the dry season in fishpond V2 with higher P concentrations. The concentrations of P in all sampled water bodies strongly correlated to concentrations of chlorophyll *a* and other photosynthetic pigments (Fig. S7), and thus may represent distinct phytoplankton bloom conditions. An RDA constrained by concentrations of photosynthetic pigments explained 8% of the total bacterial community variation, and revealed a similar separation pattern between wet and dry seasons (Fig. 4b). In this analysis, the dry season bacterial communities of fishpond V2 corresponded with higher concentrations of the cyanobacteria-related pigment zeaxanthin. Aquaculture-related estimated parameters, total fish biomass and fish-feed weight, explained only 4% of the total variation in the bacterial communities and did not reveal a clear separation between the fishponds and the untreated reservoir in an RDA ordination (Fig. S6).

In addition to the effect of environmental parameters on the microbial community structure, a strong association was identified between specific ASVs and environmental parameters. Two clusters of cyanobacterial ASVs were identified, one of which seemed to be associated with higher P concentrations and the other with higher microcystin concentrations (Fig. 5a). These cyanobacterial ASVs were mostly associated with the species *Cyanobium* PCC-6307 (15/23 ASVs), with no clear differences between the two clusters. As expected, the *Cyanobacteria* enriched ASVs corresponded to higher concentrations of cyanobacterial pigment zeaxanthin (Fig. 5b).

**Fig. 5.**
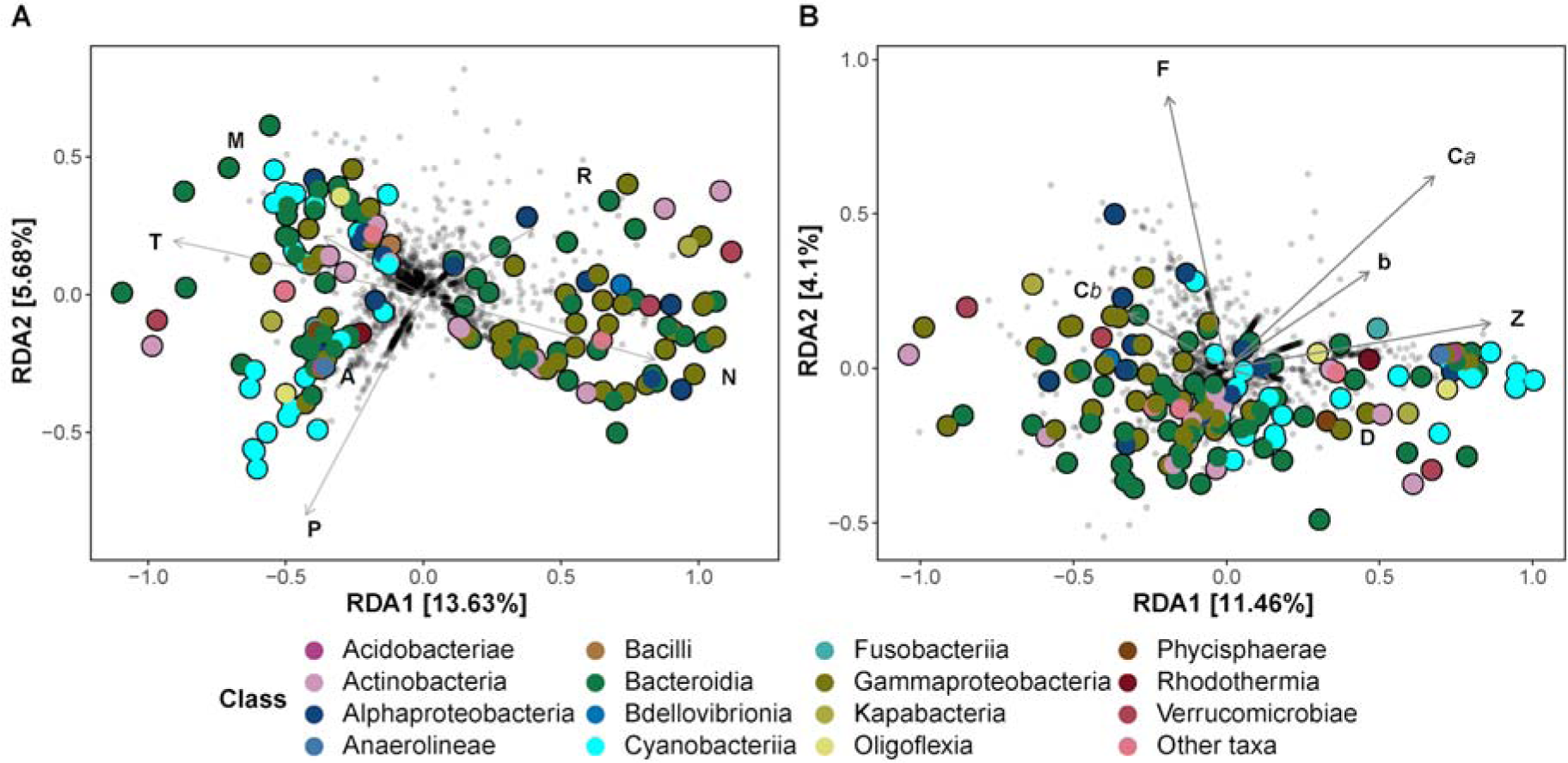
RDA ordination of bacterial ASVs constrained by physicochemical parameters (a) and photosynthetic pigments (b). The filled points represent significantly enriched ASVs, colored according to their taxonomic class. The percentage on the X and the Y axis represent the proportion of explained variance. The environmental variables are: T - temperature, N - inorganic nitrogen, P - inorganic phosphate, A - ammonia, M - microcystins, R - the N:P ratio. The pigments are: C*a*- chlorophyll *a*, C*b*- chlorophyll *b*, b - β-carotene, D - Dinoxanthin, F - fucoxanthin, Z - Zeaxanthin.

## 4. Discussion

Aquaculture facilities are anthropogenically altered hypertrophic ecosystems. Densely stocked aquaculture fishponds are supplemented with kilograms of fish-feed daily, which in most cases is not fully consumed. The remaining fish-feed, combined with fish excretions, leads to an increase in concentrations of nutrients in the water. In addition, the fishponds often receive various pharmaceuticals (Bricknell and Dalmo, 2005; Gudding and Van Muiswinkel, 2013; Sommerset et al., 2005) including antibiotics (Cháfer-Pericás et al., 2010; Kirchhelle, 2018; Lalumera et al., 2004). The resulting conditions, which are often hypereutrophic, are very different from natural freshwater environments (Klippel et al., 2020). At our study site, we examined two intensive aquaculture fishponds that are connected to an operational reservoir, which does not contain fish biomass and is not subjected directly to the intense aquaculture practices. At the initiation of this study, we expected the water quality in the fishponds, as well as the fishpond microbiomes, to differ significantly from those in the more pristine reservoir. Indeed, the nutrient concentrations in the fishponds were different from those in the reservoir, characterized by higher P and lower N concentrations, leading to differences in the N:P ratio. Therefore, it was surprising that the bacterial communities of the three water bodies were not very different, and their dynamics were primarily dictated, somewhat predictably, by seasonal (non- anthropogenic) parameters. Similar dynamics were observed recently in the Lake Taihu, where seasonality was shown to “overwhelm” net-pens aquaculture activity (Zeng et al., 2019). Our results extend this observation to a system where the water bodies are separate (linked by limited water flow), and differences in water quality (nutrients, chl-*a*) between the water bodies are much more pronounced.

### 4.1 Temperature and rainwater runoff as potential seasonal drivers of water quality and bacterial diversity

What are the potential environmental drivers of the observed seasonal patterns? We identified temperature as a significant explanatory variable of bacterial community composition, and indeed several studies have highlighted the importance of temperature as a driver of microbiome composition in aquatic environments (McFall- Ngai et al., 2013; Sunagawa et al., 2015). This includes aquaculture facilities in Lake Taihu as well as a semi-intensive aquaculture systems for growing sea bass and sea bream in Ria de Aveiro estuarine lagoon, Portugal (Duarte et al., 2019; Zeng et al., 2019). Temperature can also impact the gut microbiomes of fish and other aquatic organisms, such as tadpoles, which could also affect the structure of the fishpond microbiome (Kohl and Yahn, 2016; Kokou et al., 2018).

An additional seasonal factor that could affect water quality and subsequently the aquatic microbiome, is rainfall. During significant rain events, DARU receives runoff from the surrounding agricultural fields, carrying sediments, nutrients, fertilizers, pesticides and other soil-based compounds (Kraemer et al., 2020; Mallin et al., 2009; Maltby et al., 1995; Marmen et al., 2020; Page et al., 1995). While we lack a quantitative budget of the water and nutrient inputs into the reservoir and fishponds, our results suggest that such runoff may be important at DARU. Importantly, rainwater collected through runoff is often used as a source of water for aquaculture, and the availability of such water is likely to be reduced due to climate change (Ahmed et al., 2018). Our results highlight the need for understanding rainwater- driven links between aquaculture facilities and their surrounding terrestrial environments, through input of microorganisms, nutrients and potential contaminants (e.g., fertilizers and pesticides).

### 4.2 Aquaculture practices may affect the bacterial diversity via nutrient concentrations and N:P ratios

Despite the importance of seasonality in determining the microbial population at DARU, our results also revealed differences in the water quality and bacterial community composition between the two fishponds, and between them and the reservoir. The constantly full reservoir showed much higher concentrations of N, compared to the fishponds that are emptied annually and are refilled mostly with new underground water. It is therefore possible that seasonality, for example in rainwater runoff, exerts a stronger influence on the “older” waters of the reservoir. On the other hand, both fishponds revealed tenfold higher P concentrations, compared to the reservoir. The much higher concentrations of P in the fishponds are likely a result of the nutritional input into them, as the fish-feed used had an estimated 8% N (50% protein times 16% N in protein) compared to 1.5% PO_4_ (N:P ratio of 5.3). Indeed, the concentration of P in the water increased during the fish-stocking season.

The differences in the timing, amplitude, and potential source of nutrient input into the system led to pronounced differences in the N:P ratio between the three water bodies. The ratio between inorganic N and P is often used to infer the limiting nutrient for phytoplankton growth (Deutsch and Weber, 2012). Based on this logic, the very high N:P ratios in the reservoir suggest that primary production of this freshwater body is limited by availability of P. Fishpond D1 was likely also P-limited, whereas fishpond V2 was likely mostly N limited. The differences in nutrient concentrations, and potentially in the identity of the limiting nutrient, may be due with the aquaculture-related mismatch of the similar fish-feed input despite the severalfold lower fish biomass in fishpond V2, compared to fishpond D1, which was especially pronounced in the seasons of 2013. Notably, chl-*a* concentrations were typically higher in fishpond V2 compared to D1 and the reservoir. It is tempting to speculate that the higher chl-*a* concentrations in fishpond V2 are a result of the surplus P in this water body, although other causes such as differences in zooplankton or fish grazing cannot be ruled out. The bacterial communities of fishpond V2 during the dry season of 2013, where chl-*a* was high, also showed a strong association with higher concentrations of P. Therefore, we hypothesize that characteristics (e.g., magnitude) of the phytoplankton bloom, including also cyanobacteria, may be influenced by aquaculture-associated processes, in turn affecting the microbial communities.

### 4.3 Cyanobacterial blooms as major seasonal occurrences in aquaculture

Perhaps the most clearly observed seasonal change in the bacterial community structure were the summer cyanobacterial blooms. As a group, cyanobacteria generally exhibit optimal growth rates at relatively high temperatures, usually in excess of 25 °C (Paerl and Huisman, 2009; Robarts and Zohary, 1987) and the formation of cyanobacterial blooms is encouraged by warm and calm weather (Kanoshina et al., 2003). These conditions are prevalent during the summer months in Israel. Certain cyanobacteria can produce metabolites that cause undesirable flavors to the fish products (Lee et al., 2017; Lindholm-Lehto and Vielma, 2019; Mares et al., 2009; Paerl and Tucker, 1995; Schrader and Dennis, 2005; Tucker and Schrader, 2020), and/or various toxins that can kill both invertebrates and fish (Oberemm et al., 1999; Smith et al., 2008). Over the course of this study, cyanotoxins (microcystins or nodularins) were repeatedly observed in the particulate fraction of the reservoir and the fishponds during summer, at concentrations that, at times, were above the recommended limit for drinking water of 1 µg microcystins L^-1^ (according to guidelines set by the World Health Organization (WHO, 2019). Microcystins have repeatedly been documented in aquaculture ponds worldwide, at concentrations similar to, or higher than, observed here (Ahmed et al., 2008; Chia et al., 2009). In a survey of 485 catfish production ponds across the Southeastern USA, microcystins were detected in 47% of the ponds, with ca. 10% of the ponds containing microcystin concentrations were above the World Health Organization limit (Zimba and Grimm, 2003). These concentrations may cause damage to fish tissues, as demonstrated in a survey of fishponds in Serbia (Drobac et al., 2016). Additionally, there is evidence that microcystins affect human epidermal skin cells (Kozd ba et al., 2014), and that exposure to non-lethal doses (0.5 µg L^-1^) may contribute to the development of hepatocellular carcinoma (Pearson et al., 2010; Yoshizawa et al., 1990; Yu, 1995). Given that many aquaculture workers are often not aware of the presence of toxins in the water or their potentially harmful impact (based on personal communication with aquaculture workers around Israel), future research assessing the effects of chronic exposure, followed if needed by appropriate communication and training, are important.

While microcystins or nodularins were observed in the water of the reservoir every summer, the cyanobacteria that produced the blooms differed from year to year, and even between consecutive months of the same year (Fig. S8). Indeed, some cyanobacterial ASVs seemed to be highly specific, appearing only ephemerally, for example an ASV related to *Sphaerospermopsis torques-reginae*, which was present at high relative sequence abundance in fishpond V2 during August 2015, but was almost completely absent at other times. More generally, four main families contributed to these blooms: *Microcystaceae* (genus *Microcystis*)*, Phormidiaceae* (genus *Planktothrix*)*, Oscillatoriaceae* (an ASV related to *Planktothrocoides raciborskii*) and *Cyanobiaceae* (genus *Cyanobium*; Fig. S8). There appeared to be consistent differences between the ponds in the composition of the cyanobacterial community. For example, while the reservoir and fishpond V2 were often characterized by high relative sequence abundances of *Cyanobium*, this clade was much less common in fishpond D1, which was usually dominated by *Microcystis*.

*Microcystis* are one of the most common bloom-forming, and potentially toxin- producing, cyanobacteria (Manach et al., 2018). Blooms of *Microcystis*, including toxic species, are a well-known issue in Israel, particularly for growers of the Common carp in the northern coastal region and the Beit-Shean valley (e.g., Elkobi- Peer and Carmeli, 2015). In fact, in our previous work, we detected the presence of *Microcystis* with the potential of microcystins production in all sampled fishponds in Israel (Marmen et al., 2016). Such blooms are also common elsewhere in the world (Berdalet et al., 2018; Paerl and Paul, 2012; Zimba et al., 2001). However, in this study there were times during which, based on the 16S rRNA amplicons, *Microcystis* was almost absent, yet microcystins were present in the water. For example, a major bloom of *Planktothrocoides raciborskii* was observed in the reservoir during July 2013, with relatively high microcystin concentrations (Fig. 2, Fig. S8). However, during this bloom <1% of the 16S reads were associated with *Microcystis*. At this time, it is possible that the microcystins were produced by a *Planktothrix* that comprised ca. 8% of the 16S rRNA reads (Christiansen et al., 2003).

Finally, the cyanobacterial clade that differed between the wet and dry periods was *Cyanobium*. Relatively little is known about this clade of pico-cyanobacteria (Komárek et al., 1999; Pagels et al., 2020), despite its observed dominance in cyanobacterial blooms, including toxic ones (Li et al., 2020). They are typically found as free-living single cells, but can produce colonies in response to grazing pressure (Huber et al., 2017; Jezberová and Komárková, 2007). They have also been shown to produce allelopathic substances, although there is currently no evidence that they can produce microcystins (Costa et al., 2015; Kovács et al., 2018). Multiple ASVs of this clade were observed, and more research is needed in order to determine which factors govern the dynamics of this microorganism in the DARU system or in freshwater environments in general.

### 4.4 Are heterotrophic bacteria associated with cyanobacterial blooms or aquaculture practices?

In this study, we identified major changes in the bacterial community structure between the wet and dry seasons, and smaller changes potentially associated with the aquaculture activity. While cyanobacteria were clearly associated with the dry season, several other genera of bacteria were enriched in the dry or wet seasons. Notably, due to the fractional (relative) nature of 16S rRNA amplicon sequencing, changes in relative sequence abundance of an ASV cannot be equated with actual changes in the cell numbers of the relevant organisms (McLaren et al., 2019). The increase in cyanobacterial sequences, and the enrichment of cyanobacterial ASVs during the dry season, are likely due to blooms of these organisms, since increases were observed in both zeaxanthin, a pigment associated with cyanobacteria (Zhang et al., 2018), and cyanotoxins. For this reason, it is unclear whether the enrichment of genera such as *Polynucleobacter*, *Limnohabitans* and *Hydrogenophaga* in the wet season represents bona-fide blooms of these organisms, or whether this is a result of the increase in relative sequence abundance of cyanobacteria. In contrast, enrichment of ASVs in the dry season may be easier to interpret as bona-fide biological patterns in abundance. In this regard, the enrichment of nine ASVs belonging to the family *Saprospiracae* clade during the dry season is intriguing, as this clade is thought to be associated with the hydrolysis and utilization of complex carbon sources as well as with active predation on bacteria and algae, including *Microcystis* (Ashton and Robarts, 1987).

Additionally, both *Saprospiracae* and another group enriched in the dry season, the genus *Haliscomenobacter*, has been suggested to be associated with the presence of both cyanobacterial blooms and antibiotic resistance genes in Lake Taihu (Zhang et al., 2020a). While antibiotics are not used directly in the three sampled water bodies, they are used prophylactically at DARU at the beginning of the fish fattening cycle (beginning of dry season), including in additional fishponds connected to the system (Patil et al., 2016; Patil et al., 2020).

It has previously been suggested that the microorganisms in the aquaculture water may have a direct effect on the health of the fish (Assefa and Abunna, 2018; Kim and Lee, 2017). For example, environmental microorganisms can form the initial inocula for fish eggs or larvae, and pathogens can be transferred through the water, especially when fish biomass/density is very high. Therefore, significant effort has been invested in studying the aquatic microbiome in some aquaculture types, with the aim of using these measurements to understand fish health, to identify biomarkers/early markers of emerging diseases, or to explore the use of probiotics (Dawood et al., 2019; Kolda et al., 2020; Kuebutornye et al., 2019; Zhang et al., 2020b). In our dataset, we identify several bacterial lineages known as fish pathogens, including the genera *Aeromonas*, *Vibrio* and *Enterobacter*, in agreement with previous studies from DARU (Patil et al., 2016; Patil et al., 2020). However, due to the limited phylogenetic resolution of partially sequenced 16S rRNA gene, it is usually not possible to identify whether the observed taxa are associated specifically with pathogenic strains. For such analysis, one would need to analyze additional markers, for example genes related directly with pathogenicity. Nevertheless, we did identify one potentially interesting pattern, whereby *Vibrio*-related ASVs were found almost only in fishpond D1, and specifically during May and June of 2013 and 2014 (but not 2015), during which the *Vibrio* comprised 3-12% of the bacterial community. During these three years, fishpond D1 was used for Common carp outgrowth, however, no major cases of pathogen outbreaks were observed in the fishpond. Interestingly, during 2014 and 2015, fishpond V2 was used for outgrowth of Common carp larvae, but we haven’t detected *Vibrio* presence there. Thus, further study is needed to identify potential reasons for the strong association of *Vibrio* with a specific fishpond and time, and to what extent this is associated with changes in fish health.

## 5. Conclusions

As human impact on aquatic environments increases, it is imperative to understand how microbial communities will respond to anthropogenic stressors, and to differentiate between these responses and natural ecosystem dynamics. The aquaculture unit at DARU represents a unique semi-controlled system where water bodies under strong anthropogenic pressure – densely stocked fishponds – are tightly connected with a more “pristine” water body (the reservoir). Our results show that, even in this highly modified aquatic ecosystem, natural seasonality (e.g. temperature and rainwater runoff) is the main driver of the bacterial diversity. Perhaps the clearest manifestation of the natural cycle is the presence of cyanobacterial blooms during summer. However, the cyanobacterial bloom dynamics, which at times also included toxin-producing organisms, seem to be affected by aquaculture practices, such as fish feeding. Thus, based on the results presented here, we suggest that nutrient dynamics and possibly elemental ratios in freshwater environments, including in aquaculture, should be taken into account in the management of such ecosystems.

## Declaration of competing interest

The authors declare that they have no known competing financial interests or personal relationships that could have appeared to influence the work reported in this paper.

## Author contributions

**Sophi Marmen**: Conceptualization, Data curation, Investigation, Formal analysis, Visualization, Writing – original draft; review and editing. **Eduard Fadeev**: Data curation, Formal analysis, Visualization, Writing – original draft; review and editing. **Ashraf Al Ashhab:** Data curation, Formal analysis, Visualization, Writing – original draft; review and editing. **Ayana Benet-Perelberg:** Conceptualization, Investigation, Writing –review and editing. **Alon Naor:** Investigation, Writing –review and editing. **Hemant J. Patil:** Investigation, Writing –review and editing. **Eddie Cytryn:** Conceptualization, Funding Acquisition, Writing –review and editing. **Diti Viner- Mozzini:** Investigation, Writing –review and editing. **Assaf Sukenik:** Conceptualization, Funding Acquisition, Writing –review and editing. **Maya Lalzar:** Data curation, Formal analysis, Supervision, Writing –review and editing. **Daniel Sher:** Conceptualization, Project administration, Investigation, Formal analysis, Visualization, Writing –review and editing, Supervision, Funding Acquisition.

## Acknowledgements

We thank Tal Yahav for his help in preparing the graphical abstract.

## Funding sources

This study was supported by grant number 3-10342 from the Israeli Ministry of Science and Technology (to EC, AS and DS). Further funding was provided by grant number M-2797 of the Austrian Science Fund (to EF). The sponsors had no role in the study design, data collection, analysis or interpretation, or decision to submit for publication.

## References

1. Ahmed M, Hiller S, Luckas B. Microcystis aeruginosa bloom and the occurrence of microcystins (heptapeptides hepatotoxins) from an aquaculture pond in Gazipur, Bangladesh. Turkish Journal of Fisheries and Aquatic Sciences 2008; 8: 37–41.

2. Ahmed N, Thompson S. The blue dimensions of aquaculture: A global synthesis. Science of The Total Environment 2019; 652: 851–861.

3. Ahmed N, Ward JD, Thompson S, Saint CP, Diana JS. Blue-Green Water Nexus in Aquaculture for Resilience to Climate Change. Reviews in Fisheries Science & Aquaculture 2018; 26: 139–154.

4. Alderman D, Hastings T. Antibiotic use in aquaculture: development of antibiotic resistance–potential for consumer health risks. International journal of food science & technology 1998; 33: 139–155.

5. Ashton PJ, Robarts RD. Apparent predation of Microcystis Aeruginosa Kütz. emend elenkin by a saprospira-like bacterium in a hypertrophic lake (Hartbeespoort Dam, south Africa). Journal of the Limnological Society of Southern Africa 1987; 13: 44–47.

6. Assefa A, Abunna F. Maintenance of Fish Health in Aquaculture: Review of Epidemiological Approaches for Prevention and Control of Infectious Disease of Fish. Veterinary Medicine International 2018; 2018: 5432497.

7. Bautista-de los Santos QM, Chavarria KA, Nelson KL. Understanding the impacts of intermittent supply on the drinking water microbiome. Current Opinion in Biotechnology 2019; 57: 167–174.

8. Baxaa M, Musil M, Kummel M, Hanzlík B, Pechar L. Dissolved oxygen deficits in a shallow eutrophic aquatic ecosystem (fishpond) – Sediment oxygen demand and water column respiration alternately drive the oxygen regime. Science of The Total Environment 2020.

9. Benjamini Y, Hochberg Y. Controlling the False Discovery Rate: A Practical and Powerful Approach to Multiple Testing. Journal of the Royal Statistical Society: Series B (Methodological) 1995; 57: 289–300.

10. Berdalet E, Kudela RM, Banas NS, Bresnan E, Burford MA, Davidson K, et al. GlobalHAB: Fostering International Coordination on Harmful Algal Bloom Research in Aquatic Systems. In: Glibert PM, Berdalet E, Burford MA, Pitcher GC, Zhou M, editors. Global Ecology and Oceanography of Harmful Algal Blooms. Springer International Publishing, Cham, 2018, pp. 425–447.

11. Bricknell I, Dalmo RA. The use of immunostimulants in fish larval aquaculture. Fish & shellfish immunology 2005; 19: 457–472.

12. Bruno A, Sandionigi A, Bernasconi M, Panio A, Labra M, Casiraghi M. Changes in the Drinking Water Microbiome: Effects of Water Treatments Along the Flow of Two Drinking Water Treatment Plants in a Urbanized Area, Milan (Italy). Frontiers in Microbiology 2018; 9.

13. Callahan BJ, McMurdie PJ, Rosen MJ, Han AW, Johnson AJA, Holmes SP. DADA2: High-resolution sample inference from Illumina amplicon data. Nature Methods 2016; 13: 581–583.

14. Caporaso JG, Lauber CL, Walters WA, Berg-Lyons D, Lozupone CA, Turnbaugh PJ, Fierer N and Knight R. Global patterns of 16S rRNA diversity at a depth of millions of sequences per sample. PNAS 2011; 108: 4516–4522

15. CháferPericás C, Maquieira Á, Puchades R, Company B, Miralles J, Moreno A. Multiresidue determination of antibiotics in aquaculture fish samples by HPLC–MS/MS. Aquaculture Research 2010; 41: e217–e225.

16. Chia A, Oniye S, Ladan Z, Lado Z, Pila A, Inekwe V, et al. A survey for the presence of microcystins in aquaculture ponds in Zaria, Northern-Nigeria: Possible public health implication. African Journal of Biotechnology 2009; 8.

17. Christiansen G, Fastner J, Erhard M, Börner T and Dittmann E. Microcystin Biosynthesis in Planktothrix: Genes, Evolution, and Manipulation. Journal of Bacteriology 2003; 185: 564–572

18. Costa MS, Costa M, Ramos V, Leão PN, Barreiro A, Vasconcelos V, et al. Picocyanobacteria from a clade of marine Cyanobium revealed bioactive potential against microalgae, bacteria, and marine invertebrates. J Toxicol Environ Health A 2015; 78: 432–42.

19. Dawood MAO, Koshio S, Abdel-Daim MM, Van Doan H. Probiotic application for sustainable aquaculture. Reviews in Aquaculture 2019; 11: 907–924.

20. Deutsch C, Weber T. Nutrient ratios as a tracer and driver of ocean biogeochemistry. Annual Review of Marine Science 2012; 4: 113–141.

21. Dokulil MT, Teubner K. Cyanobacterial dominance in lakes. Hydrobiologia 2000; 438: 1–12.

22. Drobac D, Tokodi N, Lujić J, Marinović Z, Subakov-Simić G, Duli T, Važi T, Nybom S, Meriluoto J, Codd GA and Svir ev Z. Cyanobacteria and cyanotoxins in fishponds and their effects on fish tissue. Harmful Algae 2016; 55: 66–76

23. Duarte LN, Coelho FJ, Cleary DF, Bonifácio D, Martins P, Gomes NC. Bacterial and microeukaryotic plankton communities in a semi-intensive aquaculture system of sea bass (Dicentrarchus labrax): A seasonal survey. Aquaculture 2019; 503: 59–69.

24. Dusa A. Draw Venn Diagrams, 2020. https://CRAN.R-project.org/package=venn

25. Elkobi-Peer S, Carmeli S. New prenylated aeruginosin, microphycin, anabaenopeptin and micropeptin analogues from a Microcystis bloom material collected in Kibbutz Kfar Blum, Israel. Marine drugs 2015; 13: 2347–2375.

26. Garlock T, Asche F, Anderson J, Bjørndal T, Kumar G, Lorenzen K, et al. A Global Blue Revolution: Aquaculture Growth Across Regions, Species, and Countries. Reviews in Fisheries Science & Aquaculture 2020; 28: 107–116.

27. Giatsis C, Sipkema D, Ramiro-Garcia J, Bacanu GM, Abernathy J, Verreth J, et al. Probiotic legacy effects on gut microbial assembly in tilapia larvae. Scientific Reports 2016; 6: 33965.

28. Glibert PM, Landsberg JH, Evans JJ, Al-Sarawi MA, Faraj M, Al-Jarallah MA, et al. A fish kill of massive proportion in Kuwait Bay, Arabian Gulf, 2001: the roles of bacterial disease, harmful algae, and eutrophication. Harmful Algae 2002; 1: 215–231.

29. Gowen RJ. Managing eutrophication associated with aquaculture development. Journal of Applied Ichthyology 1994; 10: 242–257.

30. Grasshoff K, Kremling K, Ehrhardt M. Methods of seawater analysis: John Wiley & Sons, 2009.

31. Grze kowiak Ł, Collado MC, Salminen S. Evaluation of aggregation abilities between commensal fish bacteria and pathogens. Aquaculture 2012; 356-357: 412–414.

32. Gudding R, Van Muiswinkel WB. A history of fish vaccination: science-based disease prevention in aquaculture. Fish & shellfish immunology 2013; 35: 1683–1688.

33. Hach PF, Marchant HK, Krupke A, Riedel T, Meier DV, Lavik G, et al. Rapid microbial diversification of dissolved organic matter in oceanic surface waters leads to carbon sequestration. Scientific Reports 2020; 10: 13025.

34. Havens KE. Cyanobacteria blooms: effects on aquatic ecosystems. Springer, 2008, pp. 733–747.

35. Hooker SB, McClain CR. The calibration and validation of SeaWiFS data. Progress in Oceanography 2000; 45: 427–465.

36. Hsieh T, Ma K, Chao A. iNEXT: iNterpolation and EXTrapolation for species diversity. R package version 2.0. 19. Fecha de consulta: 13 de marzo de 2019, 2019.

37. Huber P, Diovisalvi N, Ferraro M, Metz S, Lagomarsino L, Llames ME, et al. Phenotypic plasticity in freshwater picocyanobacteria. Environ Microbiol 2017; 19: 1120–1133.

38. Jezberová J, Komárková J. Morphological transformation in a freshwater Cyanobium sp. induced by grazers. Environ Microbiol 2007; 9: 1858–62.

39. Kanoshina I, Lips U, Leppänen J-M. The influence of weather conditions (temperature and wind) on cyanobacterial bloom development in the Gulf of Finland (Baltic Sea). Harmful Algae 2003; 2: 29–41.

40. Kim JY, Lee J-L. Correlation of Total Bacterial and Vibrio spp. Populations between Fish and Water in the Aquaculture System. Frontiers in Marine Science 2017; 4.

41. Kirchhelle, C. Pharming animals: a global history of antibiotics in food production (1935–2017). Palgrave Communications 2018; 4: 1–13

42. Klippel G, Macêdo RL, Branco CWC. Comparison of different trophic state indices applied to tropical reservoirs. Lakes & Reservoirs: Science, Policy and Management for Sustainable Use 2020; 25: 214–229.

43. Kohl KD, Yahn J. Effects of environmental temperature on the gut microbial communities of tadpoles. Environmental Microbiology 2016; 18: 1561–1565.

44. Kokou F, Sasson G, Nitzan T, Doron-Faigenboim A, Harpaz S, Cnaani A, et al. Host genetic selection for cold tolerance shapes microbiome composition and modulates its response to temperature. Elife 2018; 7: e36398.

45. Kolda A, Gavrilović A, Jug-Dujaković J, Ljubešić Z, El-Matbouli M, Lillehaug A, et al. Profiling of bacterial assemblages in the marine cage farm environment, with implications on fish, human and ecosystem health. Ecological Indicators 2020; 118: 106785.

46. Komárek J, Kopecký J, Cepák V. Generic characters of the simplest cyanoprokaryotes Cyanobium, Cyanobacterium and Synechococcus. Cryptogamie Algologie 1999; 20: 209–222.

47. Kovács AW, Tóth VR, Pálffy K. The effects of interspecific interactions between bloom forming cyanobacteria and Scenedesmus quadricauda (chlorophyta) on their photophysiology. Acta Biol Hung 2018; 69: 210–223.

48. Kozd ba M, Borowczyk J, Zimol g E, Wasylewski M, Dziga D, Madeja Z, et al. Microcystin-LR affects properties of human epidermal skin cells crucial for regenerative processes. Toxicon 2014; 80: 38–46.

49. Kraemer SA, Barbosa da Costa N, Shapiro BJ, Fradette M, Huot Y, Walsh DA. A large-scale assessment of lakes reveals a pervasive signal of land use on bacterial communities. The ISME Journal 2020; 14: 3011–3023.

50. Kuebutornye FKA, Abarike ED, Lu Y. A review on the application of Bacillus as probiotics in aquaculture. Fish & Shellfish Immunology 2019; 87: 820–828.

51. Lalumera GM, Calamari D, Galli P, Castiglioni S, Crosa G, Fanelli R. Preliminary investigation on the environmental occurrence and effects of antibiotics used in aquaculture in Italy. Chemosphere 2004; 54: 661–668.

52. Li H, Barber M, Lu J, Goel R. Microbial community successions and their dynamic functions during harmful cyanobacterial blooms in a freshwater lake. Water Research 2020; 185: 116292

53. Lee J, Rai PK, Jeon YJ, Kim K-H, Kwon EE. The role of algae and cyanobacteria in the production and release of odorants in water. Environmental Pollution 2017; 227: 252–262.

54. Lindholm-Lehto PC, Vielma J. Controlling of geosmin and 2-methylisoborneol induced off-flavours in recirculating aquaculture system farmed fish—A review. Aquaculture Research 2019; 50: 9–28.

55. Liu J, Wu Y, Wu C, Muylaert K, Vyverman W, Yu H-Q, et al. Advanced nutrient removal from surface water by a consortium of attached microalgae and bacteria: A review. Bioresource Technology 2017; 241: 1127–1137.

56. Love MI, Huber W, Anders S. Moderated estimation of fold change and dispersion for RNA-seq data with DESeq2. Genome Biology 2014; 15: 550.

57. Mallin MA, Johnson VL, Ensign SH. Comparative impacts of stormwater runoff on water quality of an urban, a suburban, and a rural stream. Environmental monitoring and assessment 2009; 159: 475–491.

58. Maltby L, Forrow DM, Boxall AB, Calow P, Betton CI. The effects of motorway runoff on freshwater ecosystems: 1. Field study. Environmental Toxicology and Chemistry: An International Journal 1995; 14: 1079–1092.

59. Manach S.L, Sotton B, Huet H, Duval C, Paris A, Marie A, Yépremian C, Catherine A, Mathéron L, Vinh J, Edery M and Marie B. Physiological effects caused by microcystin-producing and non-microcystin producing Microcystis aeruginosa on medaka fish: A proteomic and metabolomic study on liver. Environmental Pollution 2018; 523–537

60. Mares J, Palikova M, Kopp R, Navratil S, Pikula J. Changes in the nutritional parameters of muscles of the common carp (Cyprinus carpio) and the silver carp (Hypophthalmichthys molitrix) following environmental exposure to cyanobacterial water bloom. Aquaculture Research 2009; 40: 148–156.

61. Marmen S, Aharonovich D, Grossowicz M, Blank L, Yacobi YZ, Sher DJ. Distribution and habitat specificity of potentially-toxic Microcystis across climate, land, and water use gradients. Frontiers in microbiology 2016; 7: 271.

62. Marmen S, Blank L, Al-Ashhab A, Malik A, Ganzert L, Lalzar M, et al. The role of land use types and water chemical properties in structuring the microbiomes of a connected lake system. Frontiers in microbiology 2020; 11: 89.

63. Martin M. Cutadapt removes adapter sequences from high-throughput sequencing reads. EMBnet.journal 2011; 10–12.

64. McFall-Ngai M, Hadfield MG, Bosch TC, Carey HV, Domazet-Lošo T, Douglas AE, et al. Animals in a bacterial world, a new imperative for the life sciences. Proceedings of the National Academy of Sciences 2013; 110: 3229–3236.

65. McLaren MR, Willis AD, Callaha BJ. Consistent and correctable bias in metagenomic sequencing experiments. eLife 2019;8:e46923.

66. McMurdie PJ, Holmes S. phyloseq: An R Package for Reproducible Interactive Analysis and Graphics of Microbiome Census Data. PLOS ONE 2013; 8: e61217.

67. Mennerat A, Nilsen F, Ebert D, Skorping A. Intensive Farming: Evolutionary Implications for Parasites and Pathogens. Evolutionary Biology 2010; 37: 59–67.

68. Muyzer G, De Waal EC and Uitierlinden G. Profiling of complex microbial populations by denaturing gradient gel electrophoresis analysis of polymerase chain reaction-amplified genes coding for 16S rRNA. Applied and Environmental Microbiology 1993; 59: 695–700

69. Nir I, Barak H, Kramarsky-Winter E, Kushmaro A. Seasonal diversity of the bacterial communities associated with petroglyphs sites from the Negev Desert, Israel. Annals of Microbiology 2019; 69: 1079–1086.

70. Oberemm A, Becker J, Codd G, Steinberg C. Effects of cyanobacterial toxins and aqueous crude extracts of cyanobacteria on the development of fish and amphibians. Environmental Toxicology: An International Journal 1999; 14: 77–88.

71. Oksanen J, Blanchet F, Friendly M, Kindt R, Legendre P, McGlinn D, et al. vegan: Community Ecology Package. R package version 2.5. 4. 2019, 2019.

72. Orland C, Emilson EJS, Basiliko N, Mykytczuk NCS, Gunn JM, Tanentzap AJ. Microbiome functioning depends on individual and interactive effects of the environment and community structure. The ISME Journal 2019; 13: 1–11.

73. Paerl H. Nutrient and other environmental controls of harmful cyanobacterial blooms along the freshwater–marine continuum. Cyanobacterial harmful algal blooms: State of the science and research needs. Springer, 2008, pp. 217–237.

74. Paerl HW, Huisman J. Climate change: a catalyst for global expansion of harmful cyanobacterial blooms. Environmental microbiology reports 2009; 1: 27–37.

75. Paerl HW, Otten TG, Kudela R. Mitigating the Expansion of Harmful Algal Blooms Across the Freshwater-to-Marine Continuum. Environmental Science & Technology 2018; 52: 5519–5529.

76. Paerl HW, Paul VJ. Climate change: Links to global expansion of harmful cyanobacteria. Water Research 2012; 46: 1349–1363.

77. Paerl HW, Tucker CS. Ecology of blue-green algae in aquaculture ponds. Journal of the World Aquaculture Society 1995; 26: 109–131.

78. Page HM, Petty RL, Meade DE. Influence of watershed runoff on nutrient dynamics in a southern California salt marsh. Estuarine, Coastal and Shelf Science 1995; 41: 163–180.

79. Pagels F, Salvaterra D, Amaro HM, Lopes G, Sousa-Pinto I, Vasconcelos V, et al. Bioactive potential of Cyanobium sp. pigment-rich extracts. Journal of Applied Phycology 2020; 32: 3031–3040.

80. Patil HJ, Benet-Perelberg A, Naor A, Smirnov M, Ofek T, Nasser A, et al. Evidence of Increased Antibiotic Resistance in Phylogenetically-Diverse Aeromonas Isolates from Semi-Intensive Fish Ponds Treated with Antibiotics. Frontiers in Microbiology 2016; 7.

81. Patil HJ, Gatica J, Zolti A, Benet-Perelberg A, Naor A, Dror B, et al. Temporal Resistome and Microbial Community Dynamics in an Intensive Aquaculture Facility with Prophylactic Antimicrobial Treatment. Microorganisms 2020; 8: 1984.

82. Pearson L, Mihali T, Moffitt M, Kellmann R, Neilan B. On the chemistry, toxicology and genetics of the cyanobacterial toxins, microcystin, nodularin, saxitoxin and cylindrospermopsin. Marine drugs 2010; 8: 1650–1680.

83. Pérez-Sánchez T, Ruiz-Zarzuela I, de Blas I, Balcázar JL. Probiotics in aquaculture: a current assessment. Reviews in Aquaculture 2014; 6: 133–146.

84. Quast C, Pruesse E, Yilmaz P, Gerken J, Schweer T, Yarza P, et al. The SILVA ribosomal RNA gene database project: improved data processing and web- based tools. Nucleic Acids Research 2012; 41: D590–D596.

85. Robarts RD, Zohary T. Temperature effects on photosynthetic capacity, respiration, and growth rates of bloom-forming cyanobacteria. New Zealand Journal of Marine and Freshwater Research 1987; 21: 391–399.

86. Rodriguez-Brito B, Li L, Wegley L, Furlan M, Angly F, Breitbart M, et al. Viral and microbial community dynamics in four aquatic environments. The ISME Journal 2010; 4: 739–751.

87. Rowe OF, Dinasquet J, Paczkowska J, Figueroa D, Riemann L, Andersson A. Major differences in dissolved organic matter characteristics and bacterial processing over an extensive brackish water gradient, the Baltic Sea. Marine Chemistry 2018; 202: 27–36.

88. Schrader KK, Dennis ME. Cyanobacteria and earthy/musty compounds found in commercial catfish (Ictalurus punctatus) ponds in the Mississippi Delta and Mississippi–Alabama Blackland Prairie. Water Research 2005; 39: 2807–2814.

89. Smith JL, Boyer GL, Zimba PV. A review of cyanobacterial odorous and bioactive metabolites: impacts and management alternatives in aquaculture. Aquaculture 2008; 280: 5–20.

90. Sommerset I, Krossøy B, Biering E, Frost P. Vaccines for fish in aquaculture. Expert review of vaccines 2005; 4: 89–101.

91. Song SK, Beck BR, Kim D, Park J, Kim J, Kim HD, et al. Prebiotics as immunostimulants in aquaculture: A review. Fish & Shellfish Immunology 2014; 40: 40–48.

92. Sunagawa S, Coelho LP, Chaffron S, Kultima JR, Labadie K, Salazar G, et al. Structure and function of the global ocean microbiome. Science 2015; 348.

93. Talbot C, Hole R. Fish diets and the control of eutrophication resulting from aquaculture. Journal of Applied Ichthyology 1994; 10: 258–270.

94. Tucker CS, Schrader KK. Off-flavors in pond-grown ictalurid catfish: Causes and management options. Journal of the World Aquaculture Society 2020; 51: 7–92.

95. Wickham H. ggplot2: elegant graphics for data analysis: springer, 2016. World Health Organization, 2019

96. Wurtsbaugh WA, Paerl HW, Dodds WK. Nutrients, eutrophication and harmful algal blooms along the freshwater to marine continuum. WIREs Water 2019; 6: e1373.

97. Yoshizawa S, Matsushima R, Watanabe MF, Harada K-i, Ichihara A, Carmichael WW, et al. Inhibition of protein phosphatases by microcystis and nodularin associated with hepatotoxicity. Journal of cancer research and clinical oncology 1990; 116: 609–614.

98. Yu SZ. Primary prevention of hepatocellular carcinoma. Journal of gastroenterology and hepatology 1995; 10: 674–682.

99. Zeng J, Lin Y, Zhao D, Huang R, Xu H, Jiao C. Seasonality overwhelms aquacultural activity in determining the composition and assembly of the bacterial community in Lake Taihu, China. Science of The Total Environment 2019; 683: 427–435.

100. Zhang Q, Zhang Z, Lu T, Peijnenburg WJGM, Gillings M, Yang X, et al. Cyanobacterial blooms contribute to the diversity of antibiotic-resistance genes in aquatic ecosystems. Communications Biology 2020a; 3: 737.

101. Zhang X, Xu X, Shen Y, Li L, Wang R, Li J. Using GFP as a biomarker to visualize the process of bacterial infection in black carp (Mylopharyngodon piceus). Aquaculture Reports 2020b; 18: 100530.

102. Zhang Y, Liu Z, Sun J, Xue C, Mao X. Biotechnological production of zeaxanthin by microorganisms. Trends in Food Science & Technology 2018; 71: 225–234.

103. Zimba PV, Grimm CC. A synoptic survey of musty/muddy odor metabolites and microcystin toxin occurrence and concentration in southeastern USA channel catfish (Ictalurus punctatus Ralfinesque) production ponds. Aquaculture 2003; 218: 81–87.

104. Zimba PV, Khoo L, Gaunt PS, Brittain S, Carmichael WW. Confirmation of catfish, Ictalurus punctatus (Rafinesque), mortality from Microcystis toxins. Journal of Fish Diseases 2001; 24: 41–47.

105. Zorz JK, Sharp C, Kleiner M, Gordon PMK, Pon RT, Dong X, et al. A shared core microbiome in soda lakes separated by large distances. Nature Communications 2019; 10: 4230

