## Supplementary information for "Seasonal dynamics are the major driver of microbial diversity and composition in intensive freshwater aquaculture"

### Figures

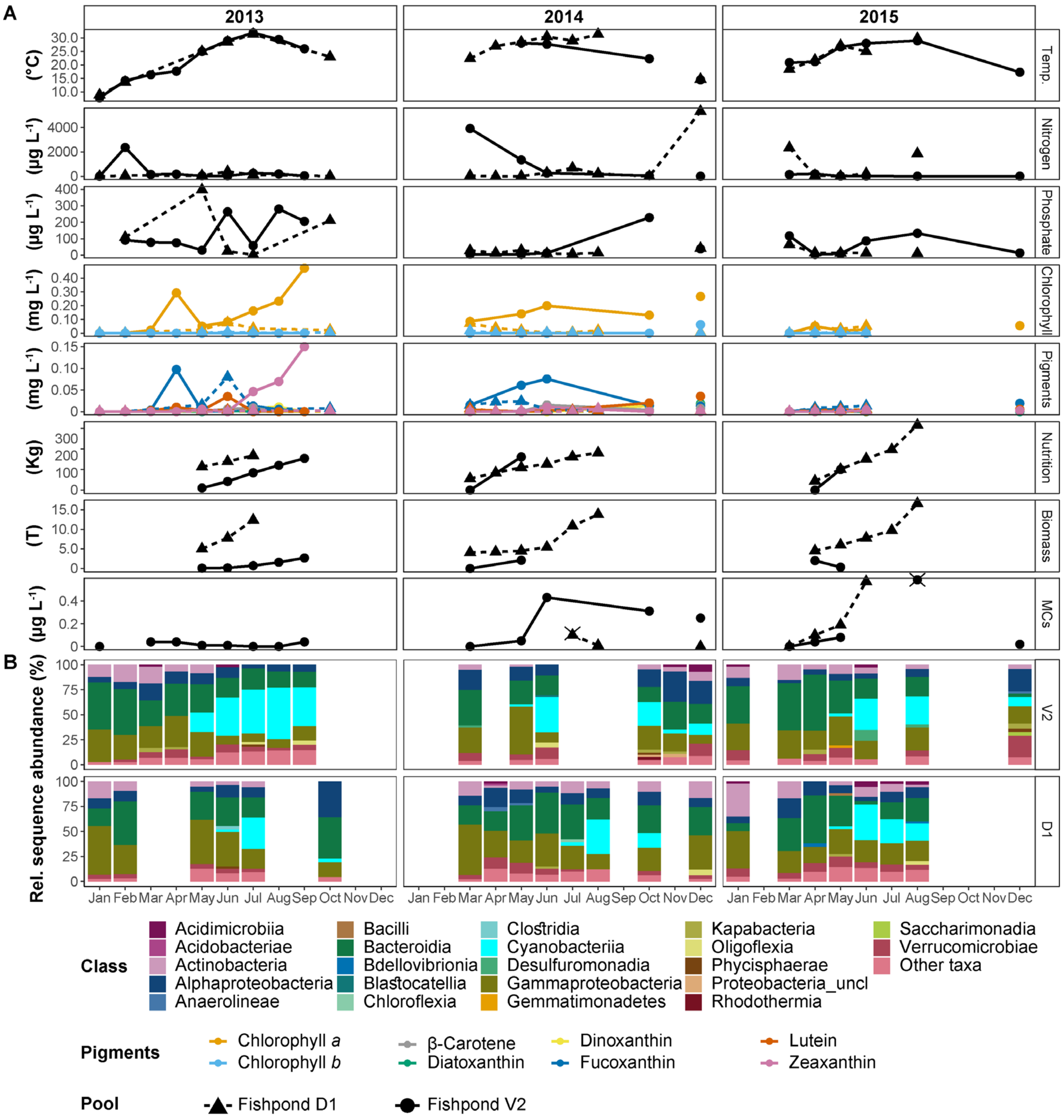


**Fig. S1.** Seasonal dynamics in the fishponds between 2013 and 2015. (a) Monthly measured physicochemical properties of the water. The different photosynthetic pigments are colored according to the legend. Shapes represent the different fishponds. Nitrogen represents the total concentration of nitrate and nitrite. In microcysteins, samples marked with '×' represent potentially underestimated concentration. (b) Sequence proportion overview of bacterial communities on a class level. The classes represented by colors according to the legend, all classes with sequence proportions below 2% were classified as "Other classes".

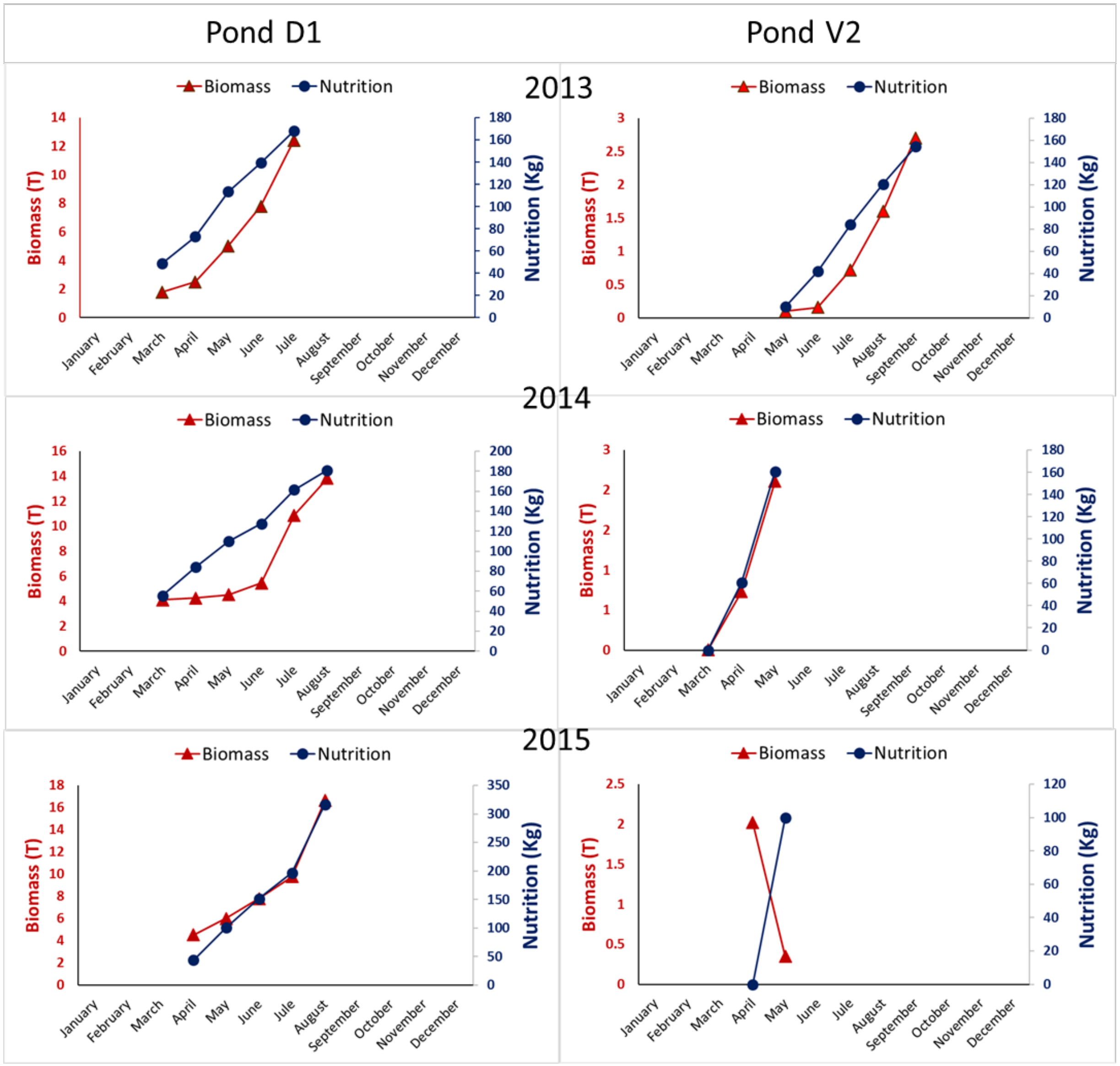


**Fig. S2.** Fish biomass and the fish-feed input differed between the two fishponds, D1 and V2. In all three seasons, the estimated fish biomass at the end of the rearing seasons in pond D1 was about fivefold higher than in the fishpond V2 (12-18 tons vs. 1.4-2.7 tons, respectively). The amounts of the fish nutrition were ca. 1.5-3% of the total biomass in pond D1 and 15-20% of the total larvae biomass in pond V2, ranged between ca. 50-350 Kg in D1 and 10-400 Kg in V2. See materials and methods for the estimation of biomass and feed from routine aquaculture measurements

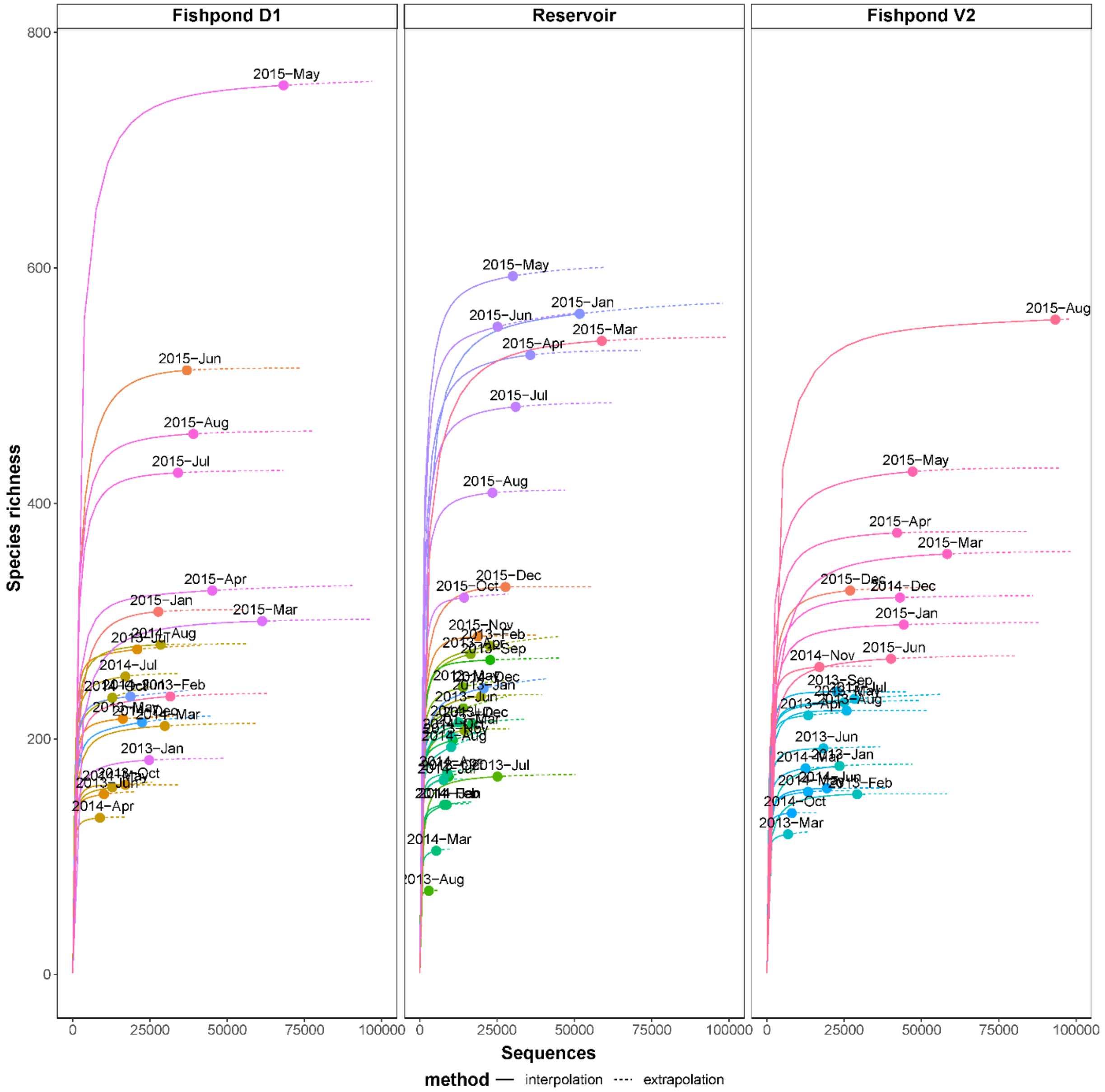


**Fig. S3.** Rarefactions of 16S rRNA gene analysis of bacterial communities in the Reservoir and the fishponds. The solid lines represent the observed accumulation with the number of reads sampled, and the dashed lines represent the extrapolated accumulation up to the double amount of reads. The observed values for each community are denoted by solid shapes. Sample-size-based rarefaction curves generated with the R-package “iNEXT”, based on the Hill number of order q = 0. The rarefaction curves for each sample were generated based on 40 equally spaced rarefied sample sizes with 100 iterations.

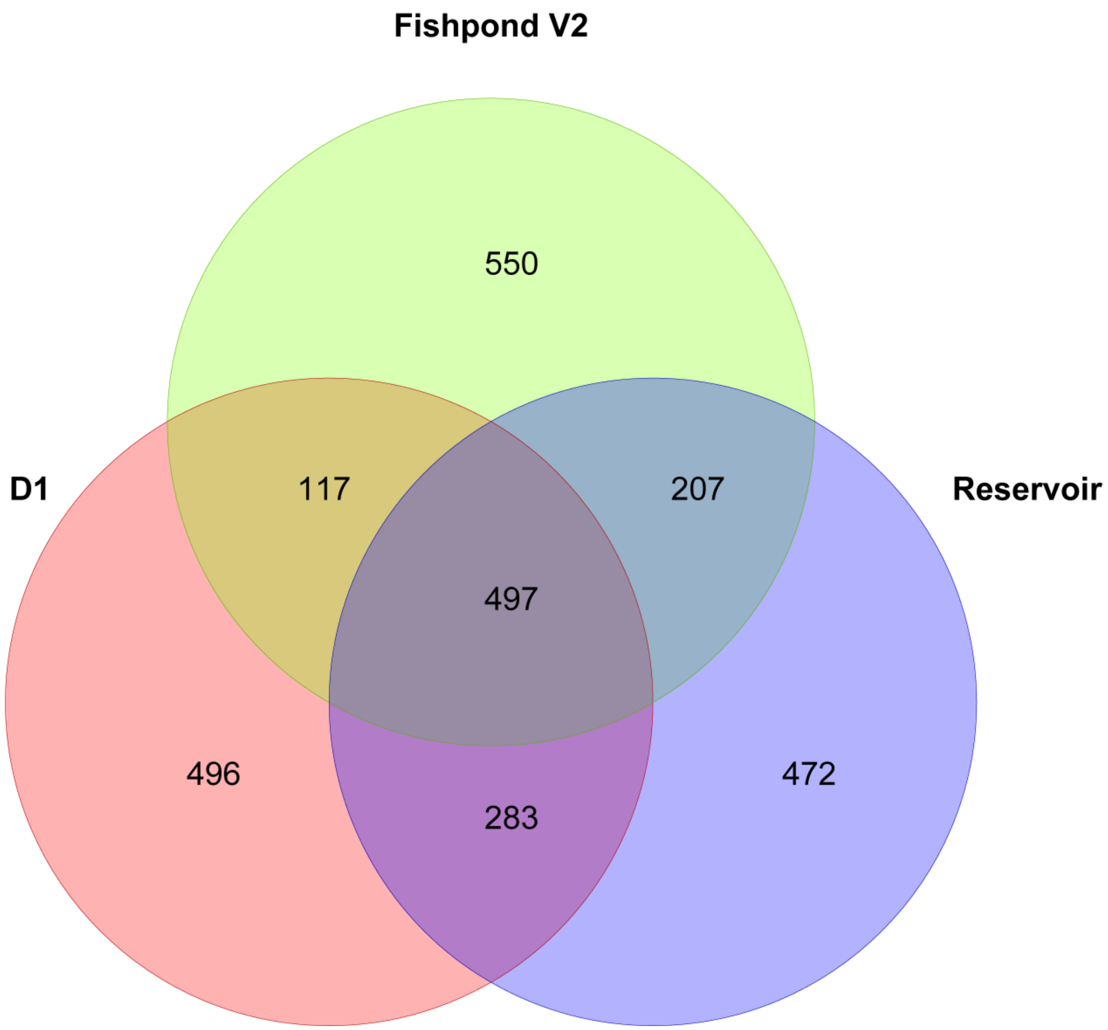


**Fig. S4.** Venn diagram of shared and unique ASVs between the bacterial communities of the Reservoir and the fishponds

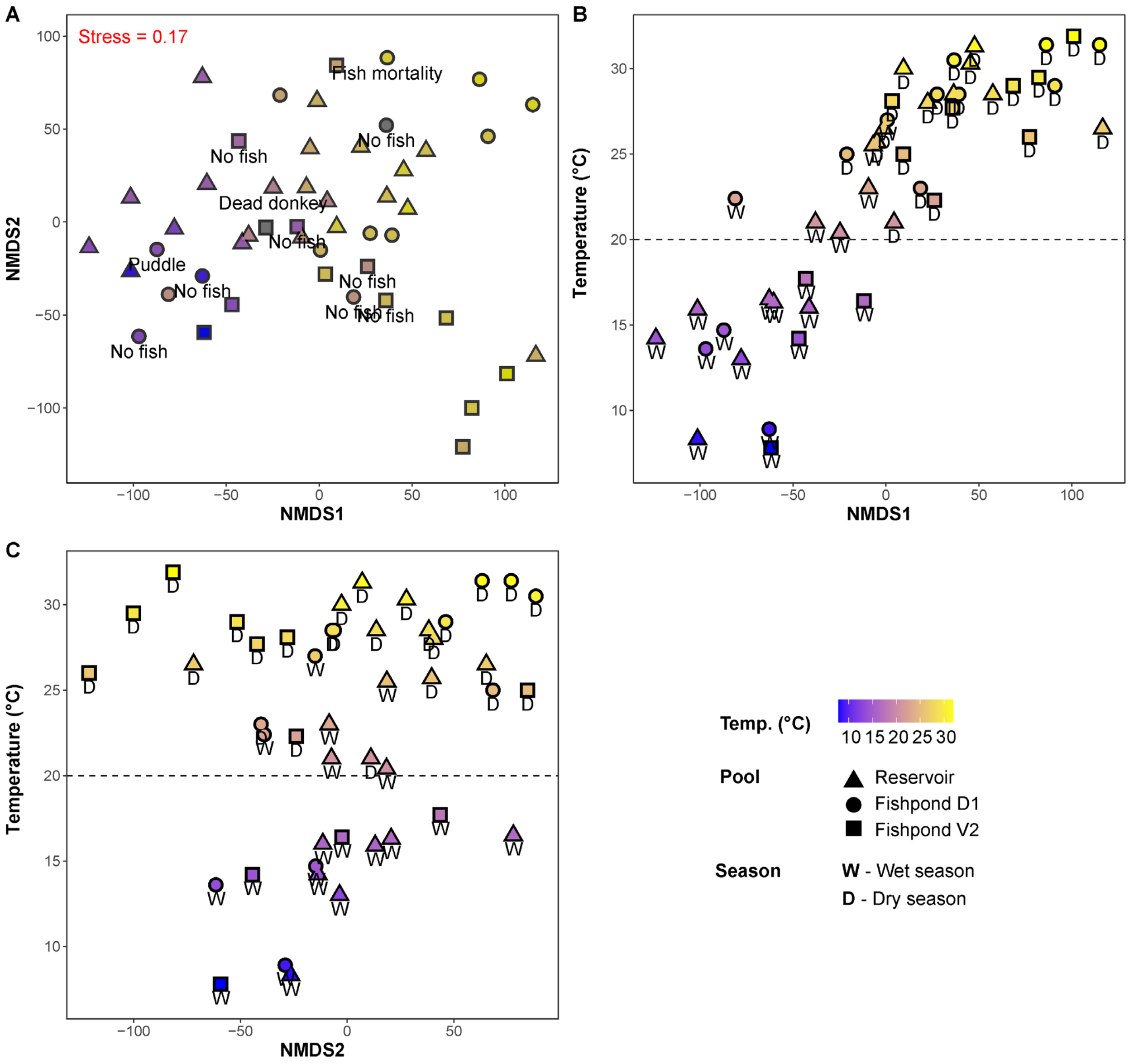


**Fig. S5.** Non-metric multidimensional scaling ordination bacterial communities of the Reservoir and the fishponds**.** The color range represents the water temperature of each sampling time point. Shapes represent the different water bodies, and letters represent the seasons, according to the legend. In panel a the labels describe special conditions with potential ecological impact. “No fish” – no fish were present in the pond at the time of sampling. “Dead donkey” – a corpse of a donkey at a stage of advanced decomposition was observed at the edge of the reservoir (donkeys were maintained at DARU to assist in minimzing plant growth around the ponds). “Puddle” – water remaining in the deepest part of the fishponds, outside of the aquaculture season.

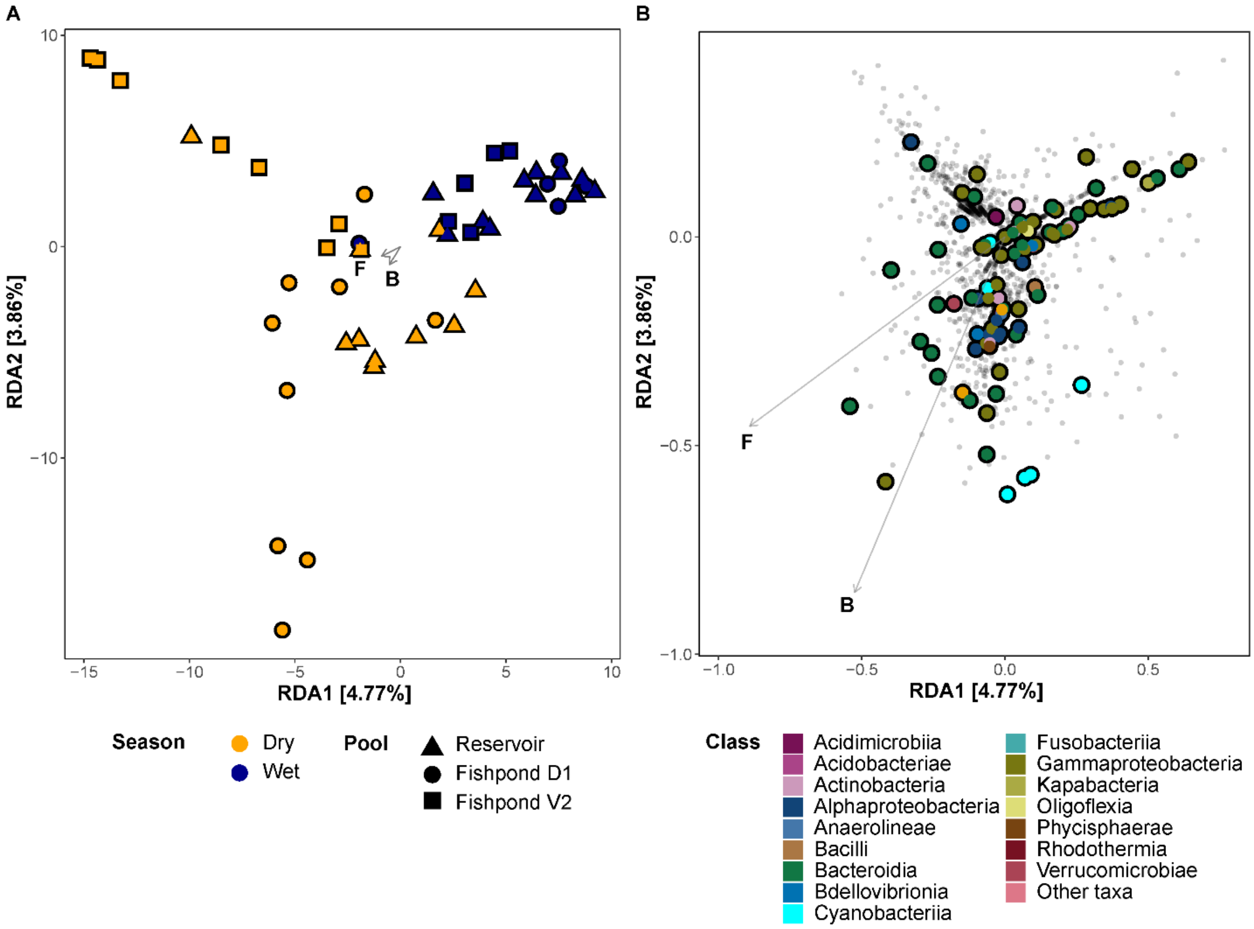


**Fig. S6.** RDA ordination of bacterial community composition constrained by aquaculture-related variables. (a) Communities constrained ordination. Colors represent the seasons and shapes represent the different pools. (b) Bacterial ASVs constrained ordination. The large points represent enriched ASVs, colored according to their taxonomic class. The environmental variables are: F - feeding input, B - total fish estimated biomass.

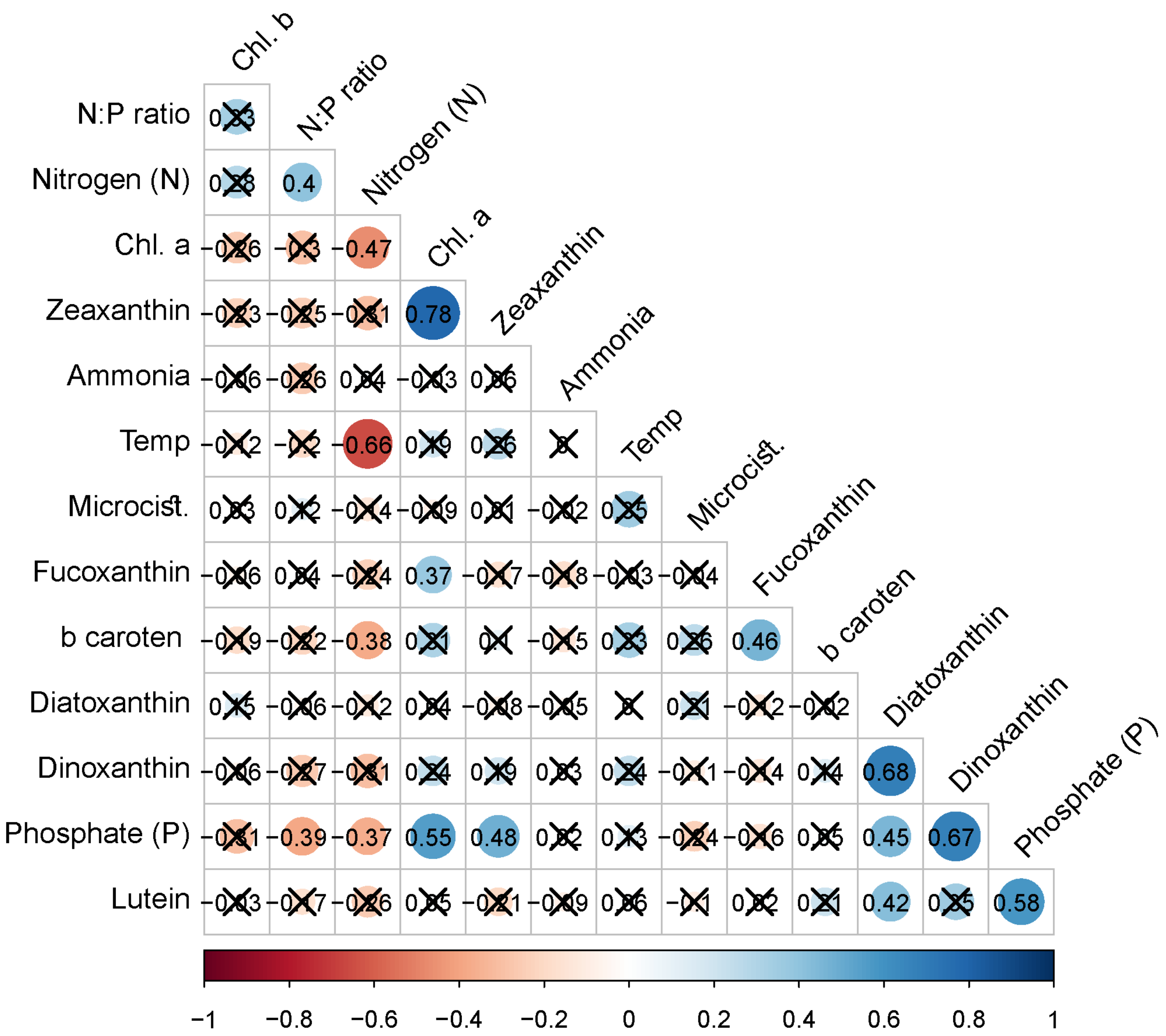


**Figure S7.** Pearsons' correlations between measured and estimated environmental parameters. The non-crossed out values represent significant correlations. Nitrogen represents the total concentration of nitrate and nitrite. The correlation strength is represented by color according to the legend. The order of the parameters is defined by hierarchical clustering of their correlations.

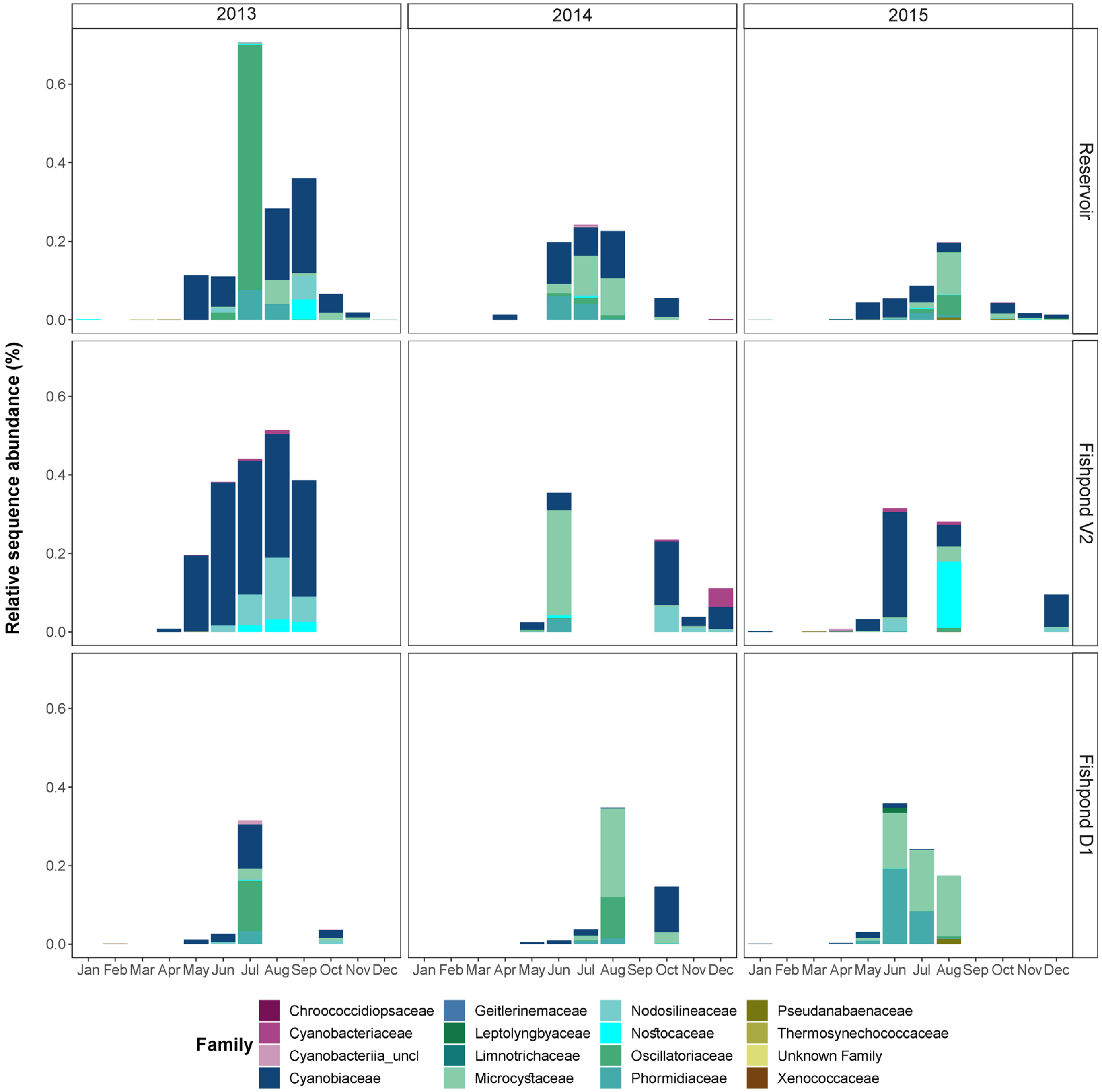


**Figure S8**: A comparison of the fraction of reads belonging to cyanobacterial families in the three water bodies over time.

#### Tables

**Table S1**: Fish culturing duration and sampling time intervals at different fishponds.

| Year | Fishpond | Fish Species | Stocking period | Number of sampled timepoints (n) | Final fish biomass [t] | Total food input [kg] |
|---|---|---|---|---|---|---|
| 2013 | D1 | *Hypophthalmichthys molitrix* | March-July | 3 | 12.4 | 421 |
| | V2 | *Cyprinus carpio* | May-September | 5 | 2.7 | 411 |
| 2014 | D1 | *Cyprinus carpio* | March-August | 6 | 13.8 | 966 |
| | V2 | *Cyprinus carpio* | April-June | 2 | 2.1 | 381 |
| 2015 | D1 | *Cyprinus carpio* | April-August | 5 | 16.6 | 721 |
| | V2 | *Cyprinus carpio* | April-May | 1 | 2 | 144 |

**Table S2**: Metadata, ASVs and diversity indices.
(provided as a separated spreadsheet)
